# Disregarding multimappers leads to biases in the functional assessment of NGS data

**DOI:** 10.1101/2023.07.04.547702

**Authors:** Michelle Almeida da Paz, Sarah Warger, Leila Taher

## Abstract

Standard ChIP-seq and RNA-seq processing pipelines typically disregard sequencing reads whose origin is ambiguous (“multimappers”). This usual practice has potentially important consequences for the functional interpretation of the data: genomic elements belonging to clusters composed of highly similar members are left unexplored. In particular, disregarding multimappers leads to the systematic underrepresentation in epigenetic studies of recently active transposons, such as AluYa5 and L1HS. Furthermore, this common strategy also has implications for transcriptomic analysis: members of repetitive gene families, such the ones including major histocompatibility complex (MHC) class I and II genes, are systematically underquantified. Based on these findings, we strongly advocate for the implementation of multimapper-aware bioinformatic genomic analyses.

## BACKGROUND

Next-generation sequencing (NGS) technologies such as Chromatin Immunoprecipitation followed by sequencing (ChIP-seq) [**1**] and RNA-seq [**2**] have emerged as the state of the art for investigating the (epi-)genome and the transcriptome. ChIP-seq and RNA-seq reads are typically short, with customary protocols recommending 1 **×** 50 bp and 2 × 75 bp, respectively [**3,4**], and such read lengths are insufficient to completely span many of the repetitive elements that abound in complex eukaryotic genomes. As a consequence, standard analysis pipelines struggle to unambiguously trace the locus from which the reads have arisen. This is well recognized by scientists working on the function and evolution of transposable elements, which have often expressed their concerns about most studies disregarding more than half of the human genome [**5–7**] and proposed several strategies to alleviate the problem. While simple strategies acknowledge ambiguously mapping reads (“multimappers”) by dividing the number of reads assigned to a locus by the total number of loci to which the reads map (e.g., [**8,9**]), more sophisticated approaches use context information to infer their origin (e.g., [**10,11**]). Standard NGS pipelines, however, habitually disregard multimappers [**12**], at most randomly reporting one of the best mappings [**13**].

With the present study we wish to draw attention to the systematic biases in downstream analysis of NGS data that result from disregarding multimappers.

## RESULTS AND DISCUSSION

Inspection of exemplary ChIP-seq ENCODE [**14**] libraries (**Additional file 1: Suppl. Table 1**) revealed that multimappers constitute a substantial proportion (9-32%) of all reads mapped to the human genome (**Fig. 1A**), although the exact numbers vary greatly depending on the mapping tool: Bwa mem [**15**] (26-32%) reported twice or more the number of multimappers than BBMap [**16**] (9-16%). As expected, a large fraction (43-79%) of multimappers mapped to regions annotated as transposable elements (TEs). Motivated by the fact that multimappers mainly originate from TEs and by the enormous expansion of repetitive TE sequences in mammalian genomes –e.g., they comprise ∼46% of the human genome–, we used ChIP-seq data to explore the impact of multimappers on the epigenetic characterisation of TEs. TE individual copies in the human genome vary widely in length (from 10 to 153,104 bp), but span a median of 231 bp, mostly reflecting the relatively recent expansion of elements from the SINE Alu family (median of 294 bp, **Fig. 1B**). Thus, the substantial fraction of multimappers derived from TEs can be explained by the fact that TE copies are not fully covered by conventional NGS reads. Specifically, we saw that while multimappers tended to be associated with evolutionary *young* TEs, such as AluYa5 and L1HS (**Additional file 1: Suppl. Table 2**), uniquely mapped reads (“unimappers”) tended to be associated with *old* TEs (p < 2.2×10^−16^, Chi-squared test; **Fig. 1C; Additional file 2: Suppl. Figs. 1-2**). This is natural, since relatively young TEs have not had enough time to accumulate variations in their sequences, but has far-reaching consequences: using standard ChIP-seq pipelines will specifically underrepresent recently active TEs, hampering their study. This situation is especially dire because TE activity is associated with diverse human diseases [**17**], and hence, the characterisation of these TEs is imperative.

**Figure 1.**
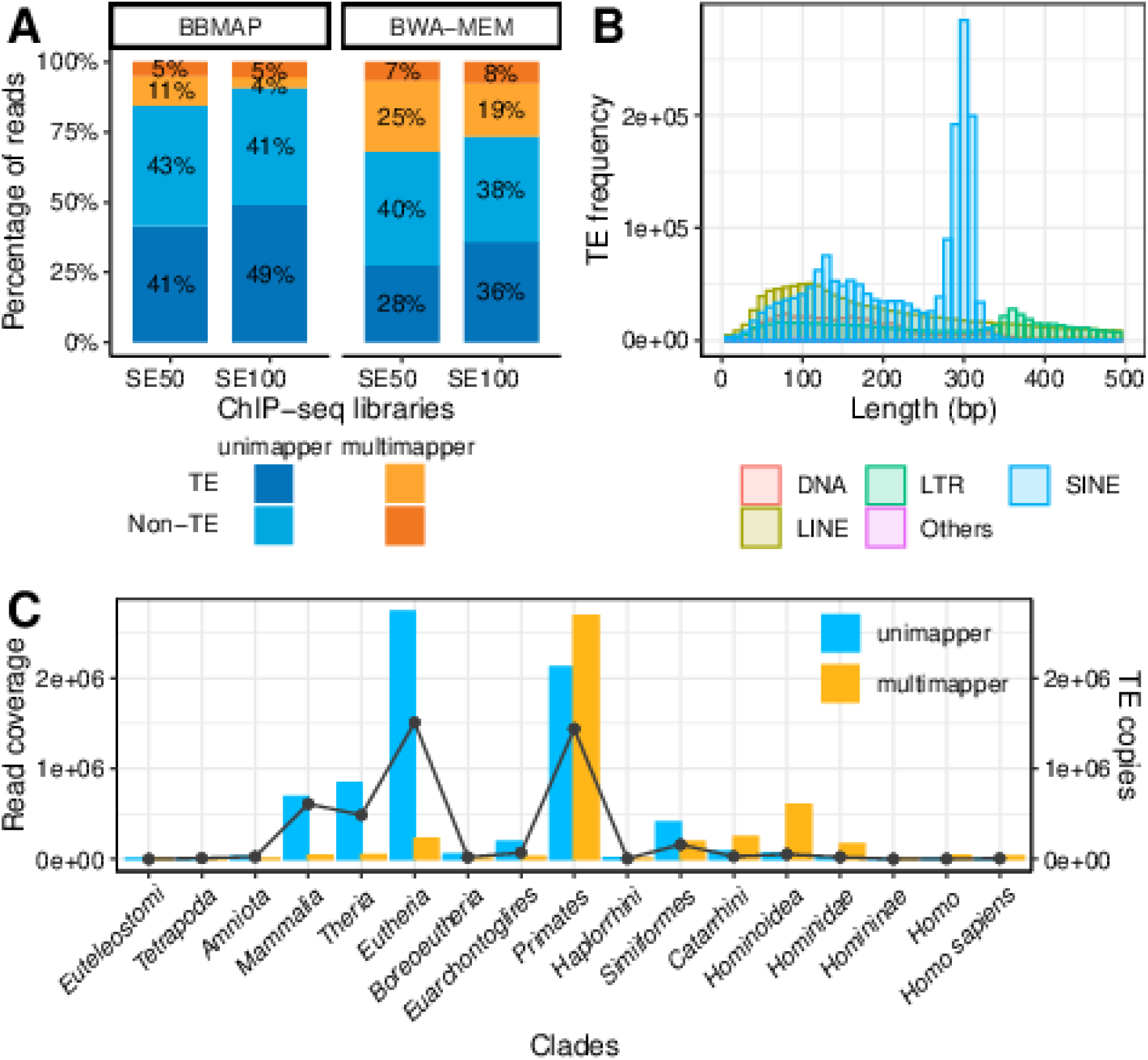
Discarding multimappers leads to epigenetic mischaracterization of young TEs. **(A)** Percentage of uni- and multimapper reads mapping to portions of the human genome annotated as TEs and not annotated as TEs (non-TE) for two ChIP-seq libraries generated by the ENCODE consortium using single-end 50bp (“SE50”) and 100bp (“SE100”) reads. Mapping was performed with two different mapping tools: with BBMap and Bwa mem. TEs are the major source of ChIP-seq multimappers in the human genome. **(B)** Length distribution of TE individual copies. Only TEs shorter than 500 bp are shown. Note that ∼74% (856 out of 1,160) of the TEs in the human genome are longer than 500 bp, spanning up to 153,104 bp. The bin width is 10 bp. TEs were classified as DNA, LTR (long terminal repeat), SINE (short interspersed nuclear element), LINE (long interspersed nuclear element) and Others (e.g., rolling-circle (RC), unknown classification). Standard NGS reads are too short to fully cover most TE copies, explaining why TEs often give rise to multimappers. **(C)** Read coverage (bar plot on the left y-axis; see Methods) per clade for the SE100 ChIP-seq library for uni- and multimappers. Reads were mapped using Bwa mem. TEs found in multiple clades (e.g., L1HS and L1P1) were assigned to the “younger” clade (*Homo* and *Hominoidea*, respectively). Only 30 out of 1,160 different TEs in the human genome (∼3%) have not been annotated to any clade and were not represented. The number of TE copies for each clade (line plot on the right y-axis) shows that the majority of TEs are *Primates* and *Eutherian*-specific. Evolutionary young TEs are prone to be underrepresented when excluding multimappers from ChIP-seq analysis.

ChIP-seq is not the only NGS technology concerned by the current prevailing approach to handling multimappers. Standard RNA-seq analysis pipelines use tools such as HTSeq-count [**18**] and STAR [**19**] for quantifying reads mapping to annotated genes, and these tools also deliberately disregard multimappers. Using RNA-seq dendritic cell libraries as an illustration of the problem, we found that ∼10% (**Fig. 2A**) and ∼5% of the fragments mapped to the human and mouse genomes, respectively, were multimappers (**Additional file 2: Suppl. Fig. 3**). Moreover, about 6% (777 out of 13,437) of the human and 4% (468 out of 12,561) of the mouse genes expressed in these cells were under-quantified by HTSeq-count and STAR geneCounts when comparing to our “multimapper-aware” strategy (**Fig. 2B; Additional file 1: Suppl. Tables 3-4; Additional file 2: Suppl. Fig. 4**). Importantly, these genes were related to specific functions intrinsic to the biology of the samples under investigation: MHC class I and II immune responses and peptide antigen binding (**Fig. 2C; Additional file 2: Suppl. Fig. 5**). In other words, disregarding multimappers may result in failure to identify relevant genes and functions. Finally, it is worth noticing that biases in quantification are likely to impact differentially expression analysis as well, since poorly expressed genes are normally filtered before testing for differential expression.

**Figure 2.**
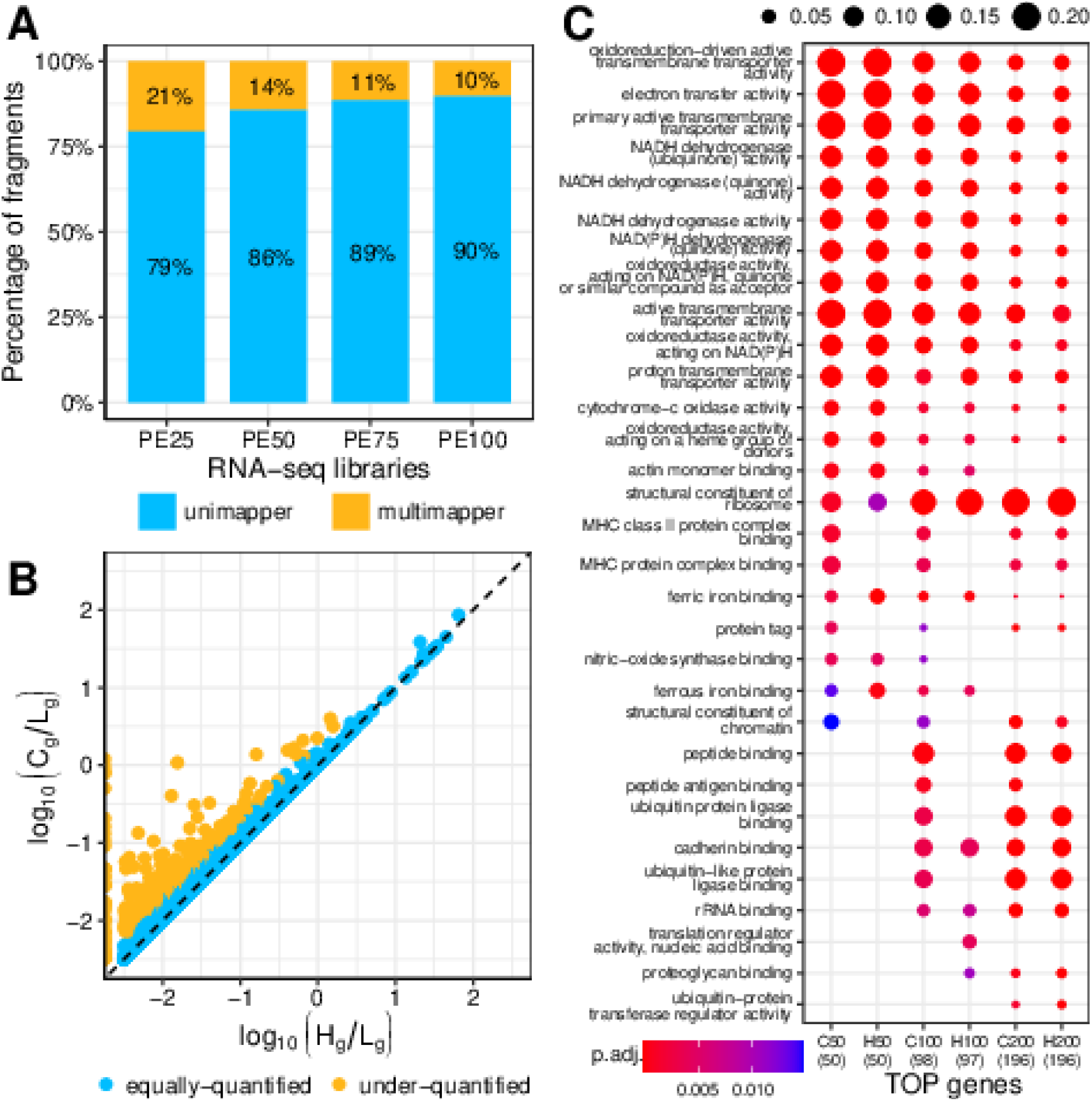
Discarding multimappers leads to functional mischaracterization of repetitive gene families. **(A)** Percentage of uni- and multimapper fragments mapping to human genome for an RNA-seq library generated by the ENCODE consortium using pair-end 100 bp (“PE100”) and thereof simulated libraries with read pairs of length 25, 50 and 75 bp (“PE25”, “PE50” and “PE75”, respectively). Within the read lengths assessed, the difference in the proportion of multimappers was modest (10-21%). **(B)** Scatter plot showing gene expression values computed with HTSeq-count using default parameters (“--nonunique none”; x-axis) and by our “multimapper-aware” strategy (y-axis) for PE100. Each dot represents a protein-coding gene and is coloured differently depending on whether it is considered (approximately) equally-quantified or under-quantified by HTSeq-count (see Methods). The dashed line indicates identical gene expression values. About 6% (777 out of 13,437) expressed genes are under-quantified when discarding multimappers. **(C)** Gene ontology (GO) enrichment analysis of the 50, 100, and 200 protein-coding genes with the highest expression values in PE100 as computed by HTSeq-count (“H50”, “H100”, and “H200”, respectively) or our “multimapper-aware” strategy (“C50”, “C100”, and “C200”, respectively). GO enrichment analysis was performed for the “molecular function” category and using the “org.Hs.eg.db” annotation for the human genome. The q-value threshold was set to 0.01. Dot size represents the ratio between the number of genes in the given GO term (y-axis) and the number of genes annotated as participating in each category (shown in brackets, below the label of each gene set on the x-axis). Dot colour indicates the *P*-value adjusted by Benjamini-Hochberg (BH, “p.adj.”). Neglecting multimappers leads to the underrepresentation of genes with specific functions.

Although, we saw little difference in the proportion of multimappers when comparing sequencing libraries with read lengths within current working standards and guidelines for ChIP-seq and RNA-seq (**Figs. 1A and 2A**), the use of substantially longer reads should improve the mappability in repetitive regions. Thus, long-read sequencing technologies (e.g., PacBio and Nanopore) are promising to minimise the effect of ambiguously mapping reads, but at higher cost and lower accuracy than NGS.

## CONCLUSIONS

In conclusion, we showed that standard NGS pipelines systematically omit genomic features containing repetitive sequences by disregarding ambiguously mapping reads (or read pairs). Furthermore, mischaracterisation of some GO functional terms is magnified when shorter reads are used. In addition, we provide a list containing the most affected features and the resulting functional consequences in our specific dataset. Finally, since the use of long-read sequencing technologies is not yet a reality in many laboratories, we raise awareness on the importance of retrieving multimappers from NGS experiments using already available strategies to gain complete understanding of the data, even if repeats are not the main focus of the analysis.

## METHODS

### Datasets

We selected four human and mouse experiments from the ENCODE Project data repository [**14**] for single-end ChIP-seq and pair-end RNA-seq with read (or read pair) length ranging from 50 to 100 bp (**Additional file 1: Suppl. Table 1**).

### Repeat annotation

Repeat annotation was obtained from the RepeatMasker track of the UCSC Genome Browser [**20**]. Immediately adjacent or overlapping annotations for TEs with the same “name” (“repName” in the RepeatMasker track) were merged. We further refer to all TEs with the same name as a TE “group”.

### Quality control and read mapping

Quality of raw ChIP-seq and RNA-seq samples was assessed using FASTQC v0.11.9 [**21**]. Reads were trimmed for adapters with Cutadapt 2.8 [**22**] and filtered with Trimmomatic v0.39 [**23**]. Bwa mem v0.7.17 [**15**] and BBMap v39.01 [**16**] were used to map reads against the human (GRCh38/hg38) and mouse (GRCm38/mm10) genome assemblies for ChIP-seq; STAR v2.7.10a [**19**] was used for RNA-seq. Gene annotations (GRCh38.p13 and GRCm38.p4) were obtained from GENCODE [**24**]. Duplicated reads were filtered out using PICARD v2.24.0 [**25**]. Reads mapping to non-chromosomal scaffolds and mitochondrial chromosome were excluded from the analysis of ChIP-seq samples. Only reads mapped in a proper pair were considered for RNA-seq data analysis; they were retrieved with SAMtools v1.10 [**26**]. The parameters used for each tool are listed in **Additional file 1: Suppl. Table 5**.

### TE group age

The oldest clade in which the TEs from a given group can be assumed to have been active was retrieved from Dfam (“Clades” column, [**27**]).

### TE group coverage

Bedmap v2.4.37 [**28**] was used to identify overlaps between the coordinates of read mappings and annotated TEs. Reads that mapped only once in the genome were considered “unimappers”; reads that mapped more than once were considered “multimappers”. Read coverage was computed for each TE group as:

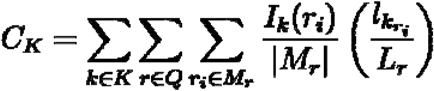

where ***K*** is the set of all copies of a TE group, ***Q*** is the set of all reads in the library, ***M***_***r***_ is the set of all loci to which read ***r*** (of ***L***_***r***_ length) mapped, and |***M***_***r***_| is the size of that set, and 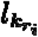 is the number of overlapping nucleotides of the ***i*** th mapping of the read ***r***_***i***_ and TE copy ***k***. For each mapping ***r***_***i***_ of ***r***

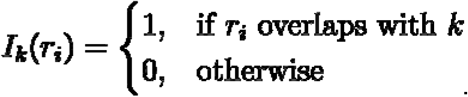

### Gene expression quantification

Multimappers were defined as read pairs (“fragments”) for which at least one read of the pair mapped more than once in the genome.

Standard gene expression quantification was performed with HTSeq-count (v2.0.2, [**18**]) using default parameters (“--nonunique none”), i.e., not accounting for multimappers. The expression value of a gene ***g*** was defined as ***H***_***g***_**/*L***_***g***_, where ***H***_***g***_ is the count for gene ***g*** assigned by HTSeq-count, and ***L***_***g***_ is the gene length as defined by its start and end coordinates in the R Ensembl BioMart database v2.54.0 [**29**].

To account for multimappers, we used a “multimapper-aware” strategy that counted fragments in genes based on the list of genes (“set S”) overlapping with the fragment mappings generated by HTSeq-count [**30**]. Specifically, gene counts were computed for each gene ***g*** as:

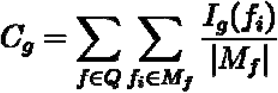

where ***Q*** is the set of all fragments in the library, ***M***_***f***_ is the set of all mappings in the transcriptome for fragment ***f*** and |***M***_***f***_| is the size of that set, and for each mapping ***f***_***i***_ of ***f***

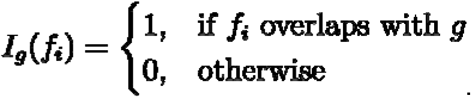

Note that if ***f***_***i***_ overlaps not only with ***g*** but also with at least another gene, then ***I***_***g***_**(*f***_***i***_**) = 0**. This is the default behaviour of HTSeq-count (**Additional file 3: Additional Material**).

A gene ***g*** was considered “expressed” if ***C***_***g***_ **> 0**. The multimapper-aware expression value of gene ***g*** was defined as ***C***_***g***_**/*L***_***g***_.

Genes were considered under-quantified by HTSeq-count if 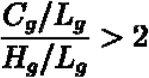, where ***H***_***g***_ is the count for gene ***g*** assigned by HTSeq-count.

Computations were repeated with simulated libraries constructed by trimming the 3’ UTR end of the read pairs to 25, 50 or 75 bp with Cutadapt 2.8 [**22**].

### Functional analysis

The 50, 100 or 200 protein-coding genes with the highest expression values were subjected to functional analysis using the “compareCluster” function of the R clusterProfiler package (v.4.6.0, [**31**]). Gene type was retrieved from R Ensembl BioMart database v2.54.0 [**29**].

## AVAILABILITY OF DATA AND MATERIALS

ChIP-seq and RNA-seq datasets were acquired, processed, and analysed from datasets described in **Additional file 1: Suppl. Table 1**. Code generated and used during the current study are available from the corresponding author on reasonable request.

## COMPETING INTERESTS

The authors declare that they have no competing interests.

## FUNDING

This work was supported by grant P33437 from the Austrian Science Fund (FWF) awarded to L.T.

## AUTHORS’ CONTRIBUTIONS

**MAP:** Methodology, Software, Visualization, Investigation, Formal analysis, Writing-Original draft preparation. **SW:** Software, Investigation. **LT:** Conceptualization, Methodology, Supervision, Writing-Reviewing and Editing.

## Acknowledgements

Not applicable.

## SUPPLEMENTARY INFORMATION

Additional file 1: Supplementary Tables 1–5.

Additional file 2: Supplementary Figures 1–5.

Additional file 3: Additional Material and Additional Figures 1–20.

## Notes

### Competing Interest Statement

The authors have declared no competing interest.

## REFERENCES

1. Robertson G, Hirst M, Bainbridge M, Bilenky M, Zhao Y, Zeng T, et al. Genome-wide profiles of STAT1 DNA association using chromatin immunoprecipitation and massively parallel sequencing. Nat Methods. 2007;4:651–7.

2. Wang Z, Gerstein M, Snyder M. RNA-Seq: a revolutionary tool for transcriptomics. Nat Rev Genet. 2009;10:57–63.

3. ENCODE Consortium. Transcription Factor ChIP-seq Data Standards and Processing Pipeline. https://www.encodeproject.org/chip-seq/transcription_factor/ (2023). Accessed 04 Apr 2023.

4. Illumina Inc. Read length recommendations. https://www.illumina.com/science/technology/next-generation-sequencing/plan-experiments/read-length.html (2023). Accessed 04 Apr 2023.

5. Lanciano S, Cristofari G. Measuring and interpreting transposable element expression. Nat Rev Genet. 2020;21:721–36.

6. O’Neill K, Brocks D, Hammell MG. Mobile genomics: tools and techniques for tackling transposons. Phil Trans R Soc B. 2020;375:20190345.

7. Teissandier A, Servant N, Barillot E, Bourc’his D. Tools and best practices for retrotransposon analysis using high-throughput sequencing data. Mobile DNA. 2019;10:52.

8. Criscione SW, Zhang Y, Thompson W, Sedivy JM, Neretti N. Transcriptional landscape of repetitive elements in normal and cancer human cells. BMC Genomics. 2014;15:583.

9. Almeida da Paz M, Taher L. T3E: a tool for characterising the epigenetic profile of transposable elements using ChIP-seq data. Mobile DNA. 2022;13:29.

10. McDermaid A, Chen X, Zhang Y, Wang C, Gu S, Xie J, et al. A New Machine Learning-Based Framework for Mapping Uncertainty Analysis in RNA-Seq Read Alignment and Gene Expression Estimation. Front Genet. 2018;9:313.

11. Yang WR, Ardeljan D, Pacyna CN, Payer LM, Burns KH. SQuIRE reveals locus-specific regulation of interspersed repeat expression. Nucleic Acids Research. 2019;47:e27–e27.

12. Landt SG, Marinov GK, Kundaje A, Kheradpour P, Pauli F, Batzoglou S, et al. ChIP-seq guidelines and practices of the ENCODE and modENCODE consortia. Genome Res. 2012;22:1813–31.

13. Deschamps-Francoeur G, Simoneau J, Scott MS. Handling multi-mapped reads in RNA-seq. Computational and Structural Biotechnology Journal. 2020;18:1569–76.

14. Davis CA, Hitz BC, Sloan CA, Chan ET, Davidson JM, Gabdank I, et al. The Encyclopedia of DNA elements (ENCODE): data portal update. Nucleic Acids Research. 2018;46:D794–801.

15. Li H, Durbin R. Fast and accurate short read alignment with Burrows-Wheeler transform. Bioinformatics. 2009;25:1754–60.

16. Bushnell B. BBMap: A Fast, Accurate, Splice-Aware Aligner. Lawrence Berkeley National Laboratory. https://escholarship.org/uc/item/1h3515gn (2014). Accessed 04 Apr 2023.

17. Hancks DC, Kazazian HH. Active human retrotransposons: variation and disease. Current Opinion in Genetics & Development. 2012;22:191–203.

18. Anders S, Pyl PT, Huber W. HTSeq—a Python framework to work with high-throughput sequencing data. Bioinformatics. 2015;31:166–9.

19. Dobin A, Davis CA, Schlesinger F, Drenkow J, Zaleski C, Jha S, et al. STAR: ultrafast universal RNA-seq aligner. Bioinformatics. 2013;29:15–21.

20. Kent WJ, Sugnet CW, Furey TS, Roskin KM, Pringle TH, Zahler AM, et al. The Human Genome Browser at UCSC. Genome Research. 2002;12:996–1006.

21. Andrews S. FastQC: A Quality Control Tool for High Throughput Sequence Data. http://www.bioinformatics.babraham.ac.uk/projects/fastqc/ (2023). Accessed 04 Apr 2023.

22. Martin M. Cutadapt removes adapter sequences from high-throughput sequencing reads. EMBnet j. 2011;17:10.

23. Bolger AM, Lohse M, Usadel B. Trimmomatic: a flexible trimmer for Illumina sequence data. Bioinformatics. 2014;30:2114–20.

24. Frankish A, Diekhans M, Ferreira AM, Johnson R, Jungreis I, Loveland J, et al. GENCODE reference annotation for the human and mouse genomes. Nucleic Acids Research. 2019;47:D766–73.

25. Broad Institute. Picard Toolkit. https://broadinstitute.github.io/picard/ (2023). Accessed 04 Apr 2023.

26. Li H, Handsaker B, Wysoker A, Fennell T, Ruan J, Homer N, et al. The Sequence Alignment/Map format and SAMtools. Bioinformatics. 2009;25:2078–9.

27. Storer J, Hubley R, Rosen J, Wheeler TJ, Smit AF. The Dfam community resource of transposable element families, sequence models, and genome annotations. Mobile DNA. 2021;12:2.

28. Neph S, Kuehn MS, Reynolds AP, Haugen E, Thurman RE, Johnson AK, et al. BEDOPS: high-performance genomic feature operations. Bioinformatics. 2012;28:1919–20.

29. Durinck S, Spellman PT, Birney E, Huber W. Mapping identifiers for the integration of genomic datasets with the R/Bioconductor package biomaRt. Nat Protoc. 2009;4:1184–91.

30. Anders S. Counting reads in features with htseq-count. https://htseq.readthedocs.io/en/release_0.11.1/count.html (2010). Accessed 04 Apr 2023.

31. Wu T, Hu E, Xu S, Chen M, Guo P, Dai Z, et al. clusterProfiler 4.0: A universal enrichment tool for interpreting omics data. The Innovation. 2021;2:100141.

