## Additional File 2 for "Disregarding multimappers leads to biases in the functional assessment of NGS data"

### Supplementary Figures

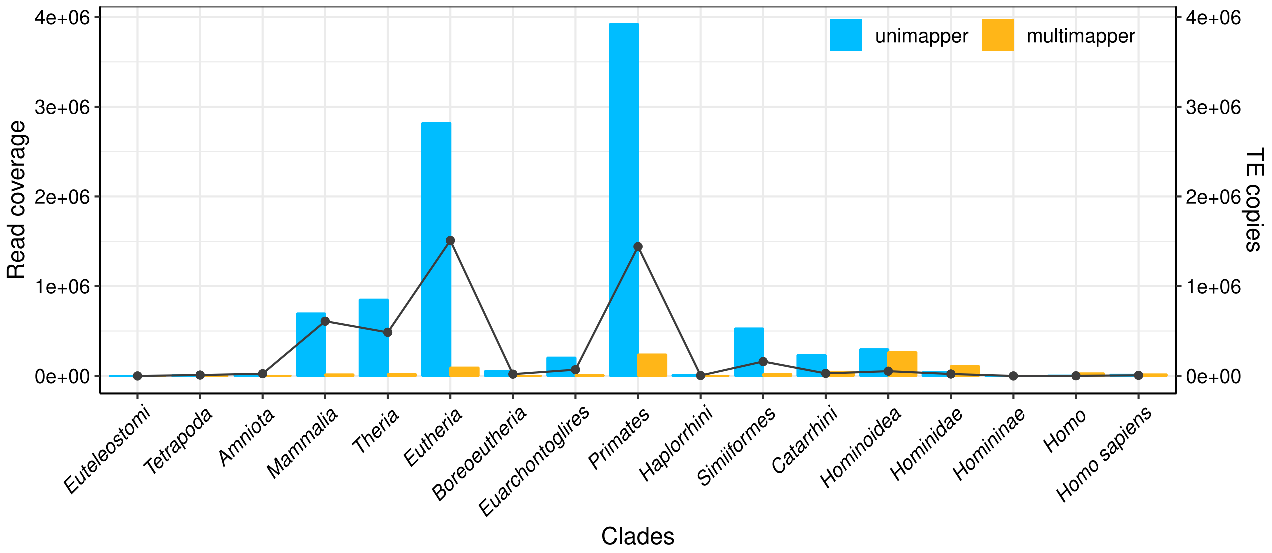


**Supplementary Figure 1. Human SE100 ChIP-seq library mapped using BBMAP.** Read coverage (left y-axis) and number of TE copies (right y-axis) per clade for uni- and multimappers. For complete description, see legend of Figure 1.

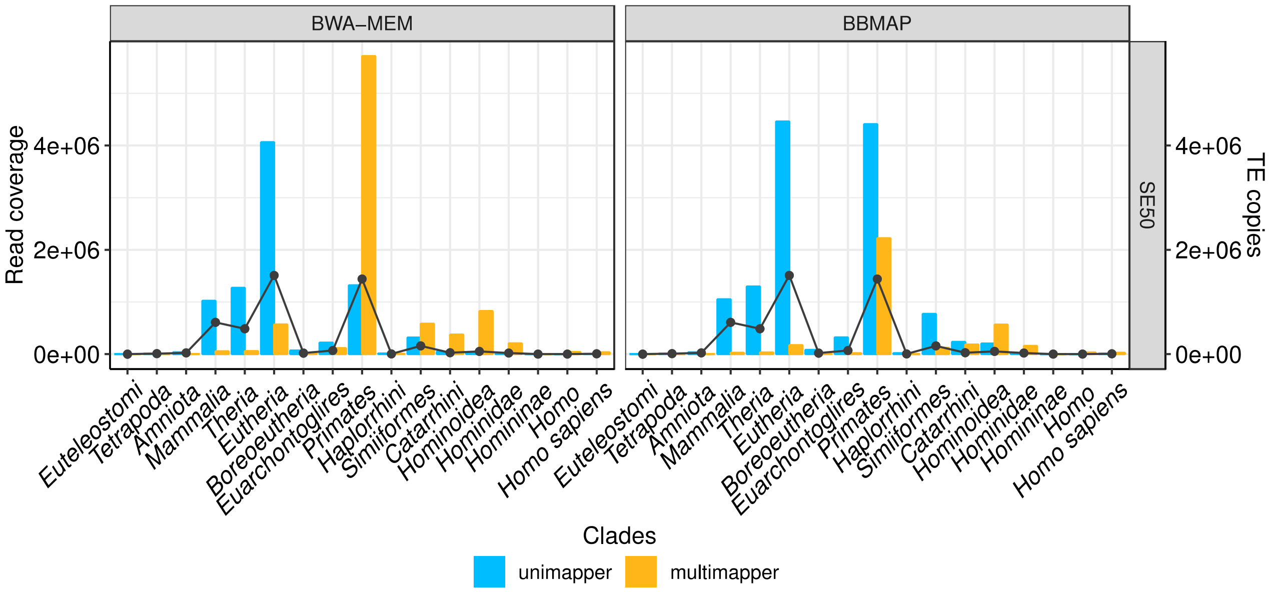


**Supplementary Figure 2. Human SE50 ChIP-seq library mapped using BWA-MEM and BBMAP.** Read coverage (left y-axis) and number of TE copies (right y-axis) per clade for uni- and multimappers. For complete description, see legend of Figure 1.

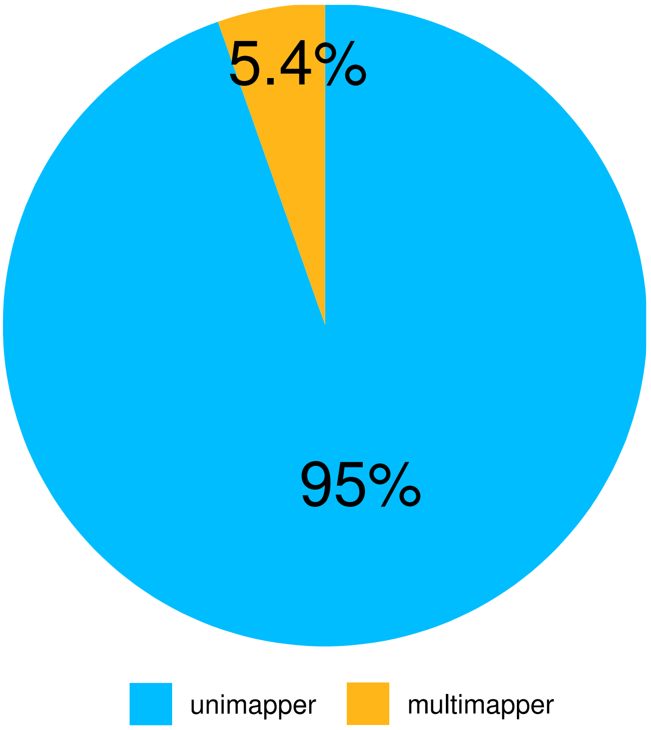


**Supplementary Figure 3. Percentage of uni- and multimapper fragments for mouse PE75 RNA-seq library.** Library was generated by the ENCODE consortium using pair-end 75 bp. Fragments were mapped to the mouse genome.

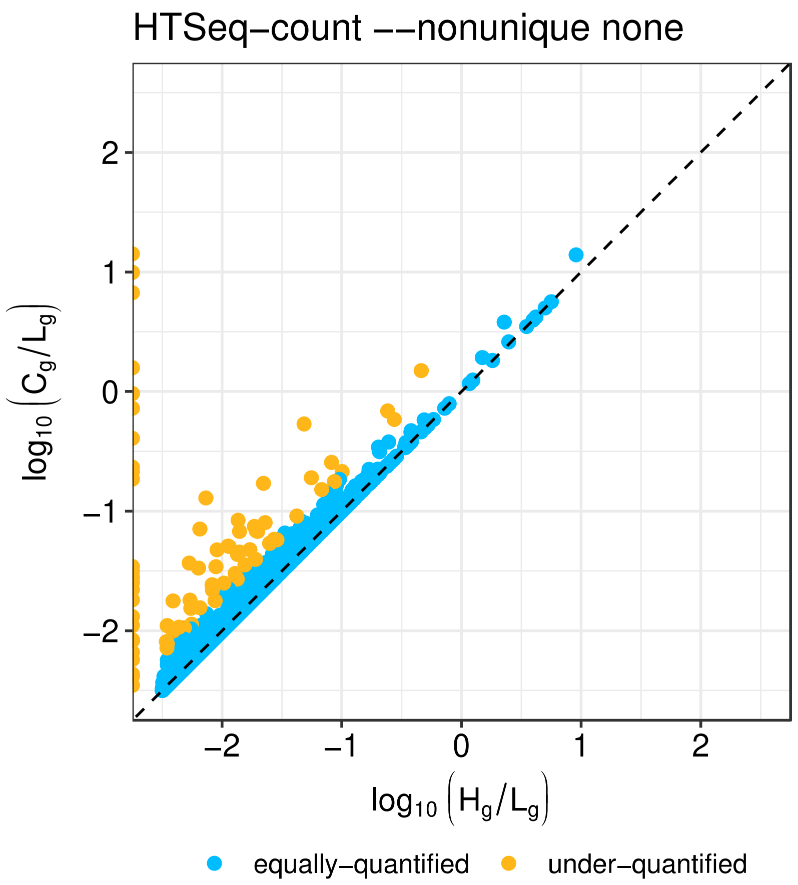


**Supplementary Figure 4. Gene under-quantification by HTSeq-count (--nonunique none) for mouse PE75 RNA-seq library.** Scatter plot showing under-quantified protein-coding genes by HTSeq-count using default parameters ("--nonunique none"; x-axis), when comparing to "multimapper-aware" expression values (y-axis). About 4% (468 out of 12,561) expressed genes are under-quantified when discarding multimappers. For complete description, see legend of Figure 2B.

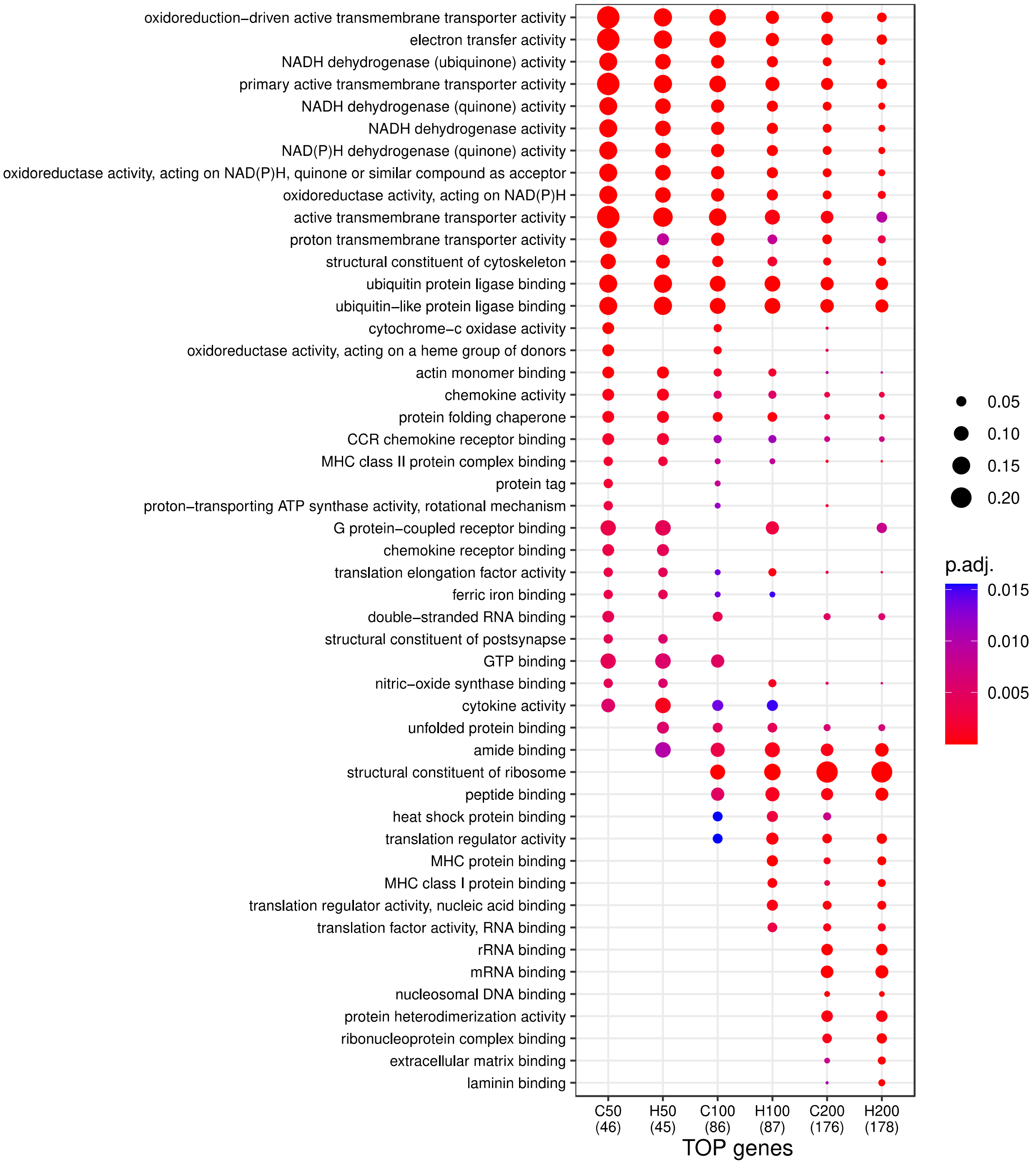


**Supplementary Figure 5. Functional underrepresentation by HTSeq-count (--nonunique none) for Mouse PE75 RNA-seq library.** Dot plot showing gene ontology (GO) enrichment analysis of the 50, 100, and 200 protein-coding genes with the highest expression values as computed by HTSeq-count using default parameters ("--nonunique none") ("H50", "H100", and "H200", respectively) or our "multimapper-aware" strategy ("C50", "C100", and "C200", respectively). For complete description, see legend of Figure 2C.
