## Additional File 3 for "Disregarding multimappers leads to biases in the functional assessment of NGS data"

### Additional Material

We observed that about 6% (777 out of 13,437) of the expressed genes were under-quantified by HTSeq-count using default parameters ("--nonunique none") for the human pair-end 100 bp ("PE100") RNA-seq library (see Methods). Here, we investigated computationally trimming of 100 bp (25, 50, 75 bp) and observed that the percentage of genes reported as under-quantified by discarding multimappers was magnified (~6.5-12%) by decreasing the read length (**Additional Figure 1**). Besides that, we evaluated other HTSeq-count "--nonunique" options. Genes were considered under-quantified if the expression value calculated using the "multimapper aware" strategy was at least two times greater than the expression value for HTSeq-count (see Methods). Genes were considered over-quantified if the expression value for HTSeq-count was at least two times greater than the expression value calculated using the "multimapper aware" strategy. Otherwise, genes were considered equally-quantified. For the modes all (**Additional Figure 2**) and fraction (**Additional Figure 3**), the percentage of over-quantified genes ranged between ~1.2-4%. For the "--nonunique random", about 1.4-4% of the expressed genes were over-quantified, and about 0.4-0.7% were under-quantified (**Additional Figure 4**).

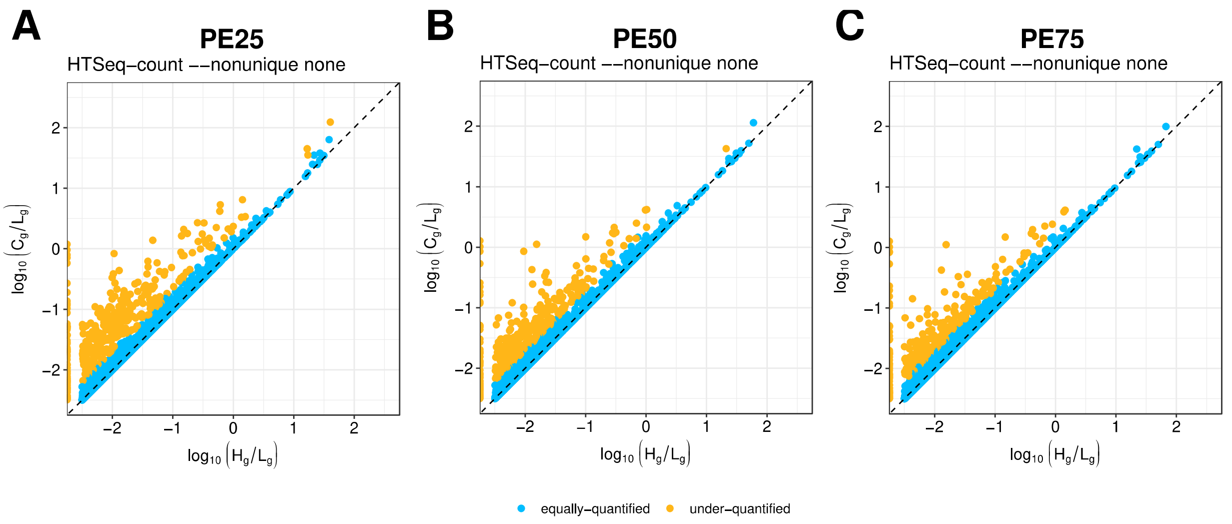


**Additional Figure 1. Gene misquantification by HTSeq-count (--nonunique none) for computational trimming of human PE100 RNA-seq library.** Scatter plot showing under-quantified protein-coding genes by HTSeq-count using default parameters ("--nonunique none"; x-axis), when comparing to "multimapper-aware" expression values (y-axis). Percentage of expressed genes that are under-quantified when discarding multimappers for **(A)** PE25: 12% (1,728 out of 14,046). **(B)** PE50: 8% (1,061 out of 13,599). **(C)** PE75: 6.5% (873 out of 13,528). For complete description, see legend of Figure 2B.

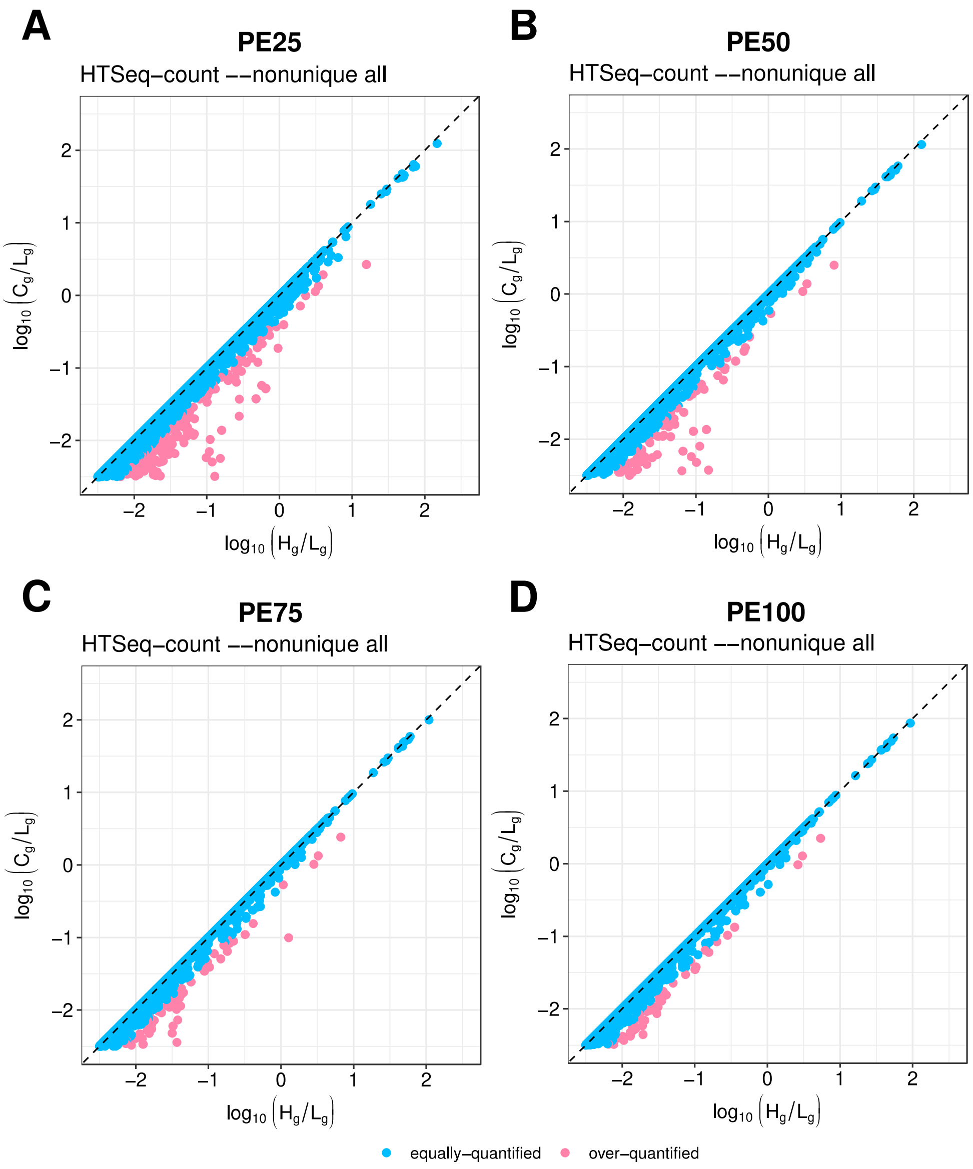


**Additional Figure 2. Gene misquantification by HTSeq-count (--nonunique all) for computational trimming of human PE100 RNA-seq library.** Scatter plot showing over-quantified protein-coding genes by HTSeq-count using parameter ("--nonunique all"; x-axis), when comparing to "multimapper-aware" expression values (y-axis). Percentage of expressed genes that are over-quantified for **(A)** PE25: 4% (589 out of 14,854). **(B)** PE50: 2% (288 out of 14,323). **(C)** PE75: 1.5% (216 out of 14,253). **(D)** PE100: 1.2% (173 out of 14,160). For complete description, see legend of Figure 2B.

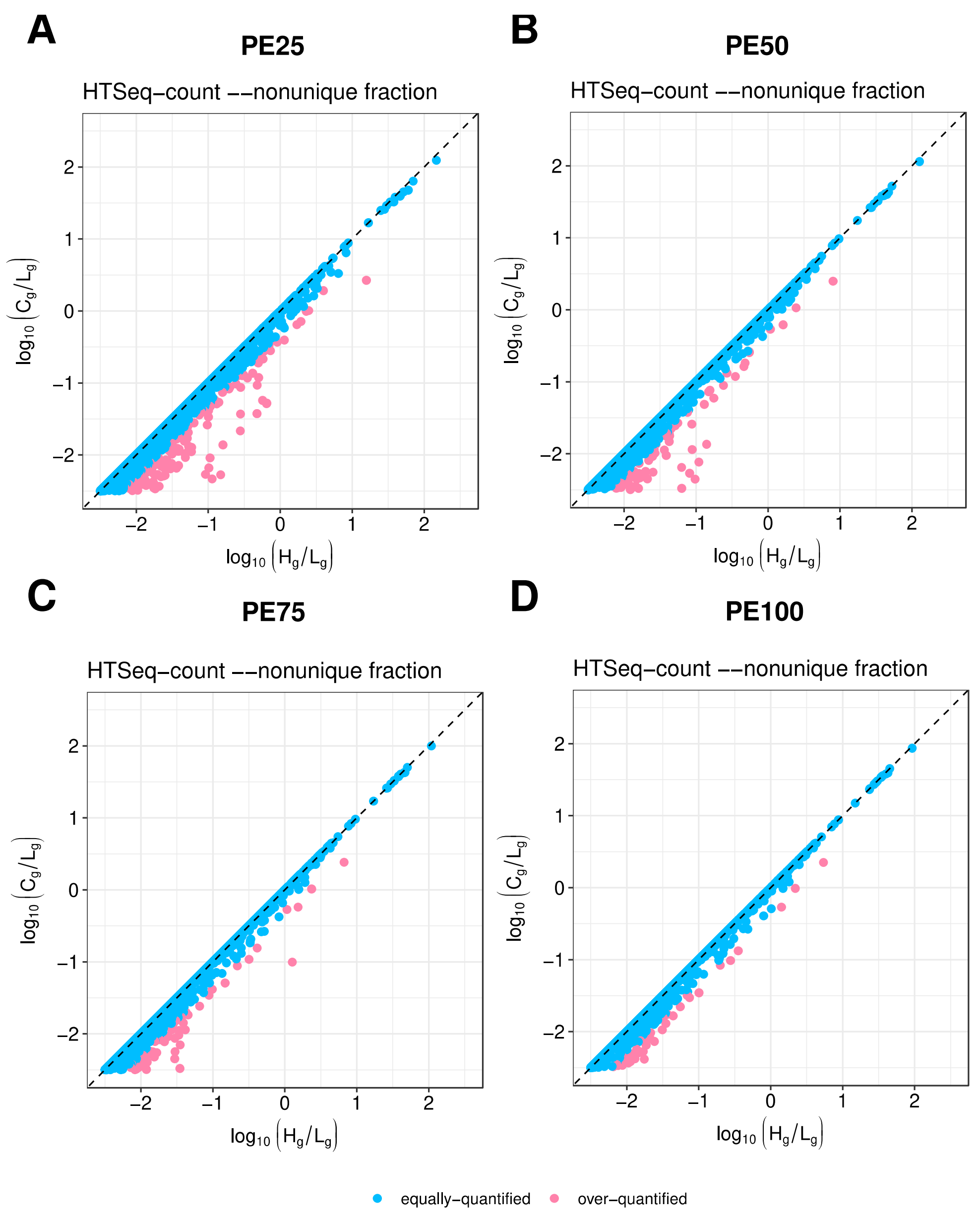


**Additional Figure 3. Gene misquantification by HTSeq-count (--nonunique fraction) for computational trimming of human PE100 RNA-seq library.** Scatter plot showing over-quantified protein-coding genes by HTSeq-count using parameter ("--nonunique fraction"; x-axis), when comparing to "multimapper-aware" expression values (y-axis). Percentage of expressed genes that are over-quantified for **(A)** PE25: 4% (577 out of 14,854). **(B)** PE50: 2% (282 out of 14,323). **(C)** PE75: 1.5% (205 out of 14,253). **(D)** PE100: 1.2% (167 out of 14,160). For complete description, see legend of Figure 2B.

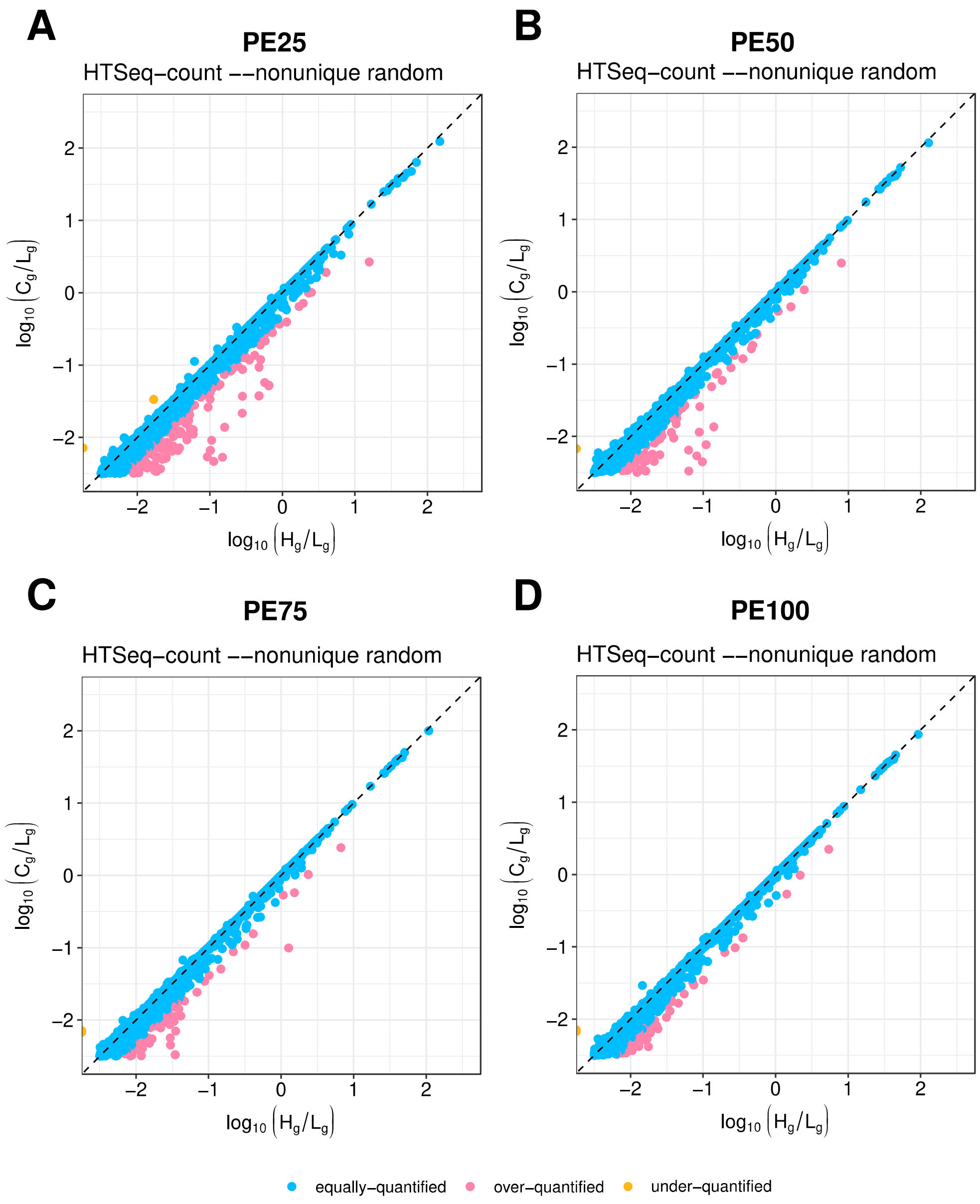


**Additional Figure 4: Gene misquantification by HTSeq-count (--nonunique random) for computational trimming of human PE100 RNA-seq library.** Scatter plot showing under and over-quantified protein-coding genes by HTSeq-count using parameter ("--nonunique random"; x-axis), when comparing to "multimapper-aware" expression values (y-axis). Percentage of expressed genes for **(A)** PE25: 4% (610 out of 14,854) are over-quantified and 0.7% (108 out of 14,854) are under-quantified. **(B)** PE50: 2% (300 out of 14,323) are over-quantified and 0.5% (70 out of 14,323) are under-quantified. **(C)** PE75: 1.6% (225 out of 14,253) are over-quantified and 0.5% (71 out of 14,253) are under-quantified. **(D)** PE100: 1.4% (192 out of 14,160) are over-quantified and 0.4% (60 out of 14,160) are under-quantified. For complete description, see legend of Figure 2B.

We showed that genes with particular functions to the studied RNA-seq samples were underrepresented when disregarding multimappers, and it was more dramatic the shorter the read length (**Additional Figure 5**). Furthermore, we evaluated other HTSeq-count "--nonunique" options ("all", "fraction" and "random") and found other misrepresented functional GO terms (**Additional Figures 6-8**).

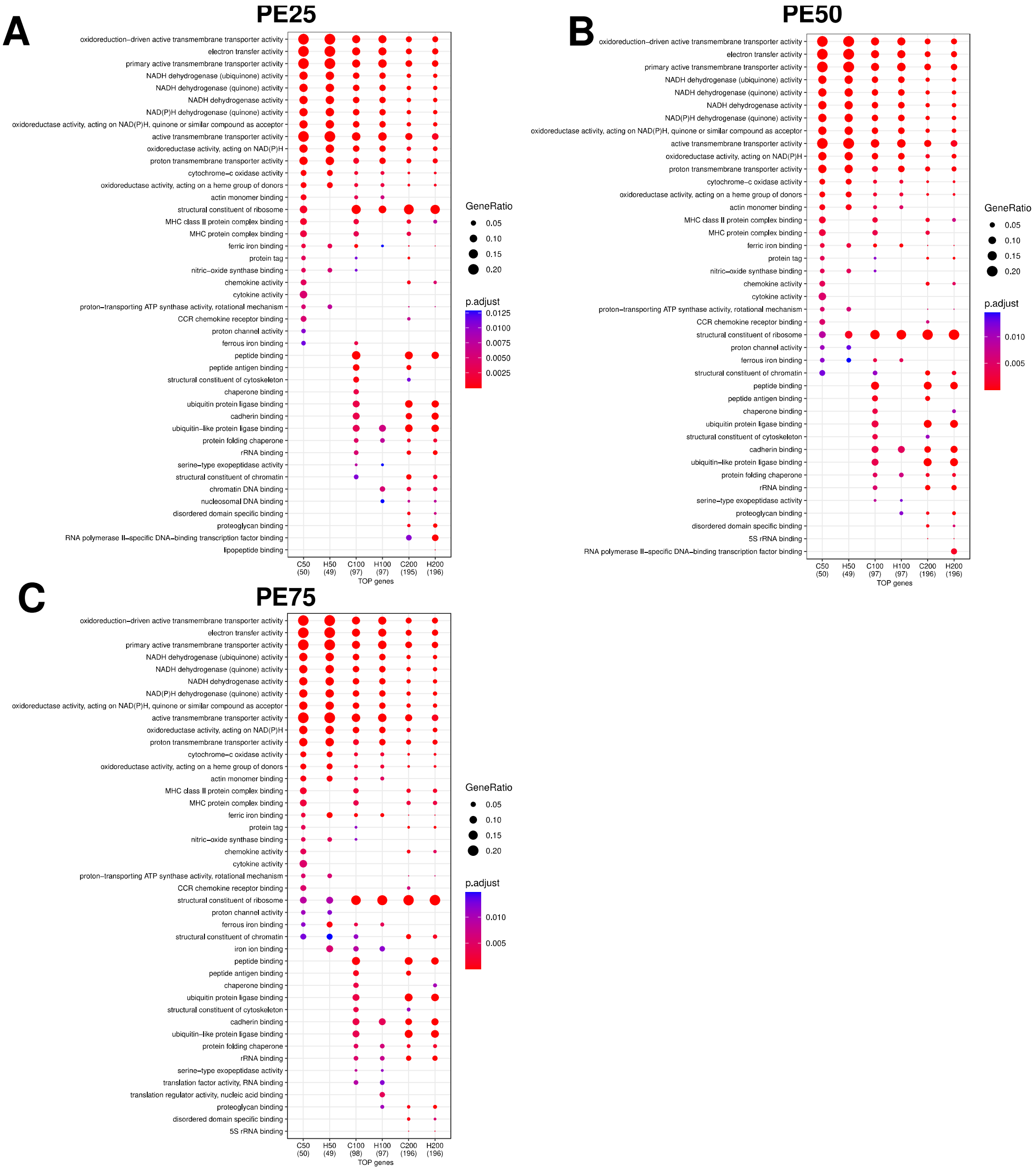


**Additional Figure 5. Functional misrepresentation by HTSeq-count (--nonunique none) for computational trimming of human PE100 RNA-seq library.** Dot plot showing gene ontology (GO) enrichment analysis of the 50, 100, and 200 protein-coding genes with the highest expression values as computed by HTSeq-count using default parameters ("--nonunique none") ("H50", "H100", and "H200", respectively) or our "multimapper-aware" strategy ("C50", "C100", and "C200", respectively). **(A)** PE25. **(B)** PE50. **(C)** PE75. For complete description, see legend of Figure 2C.

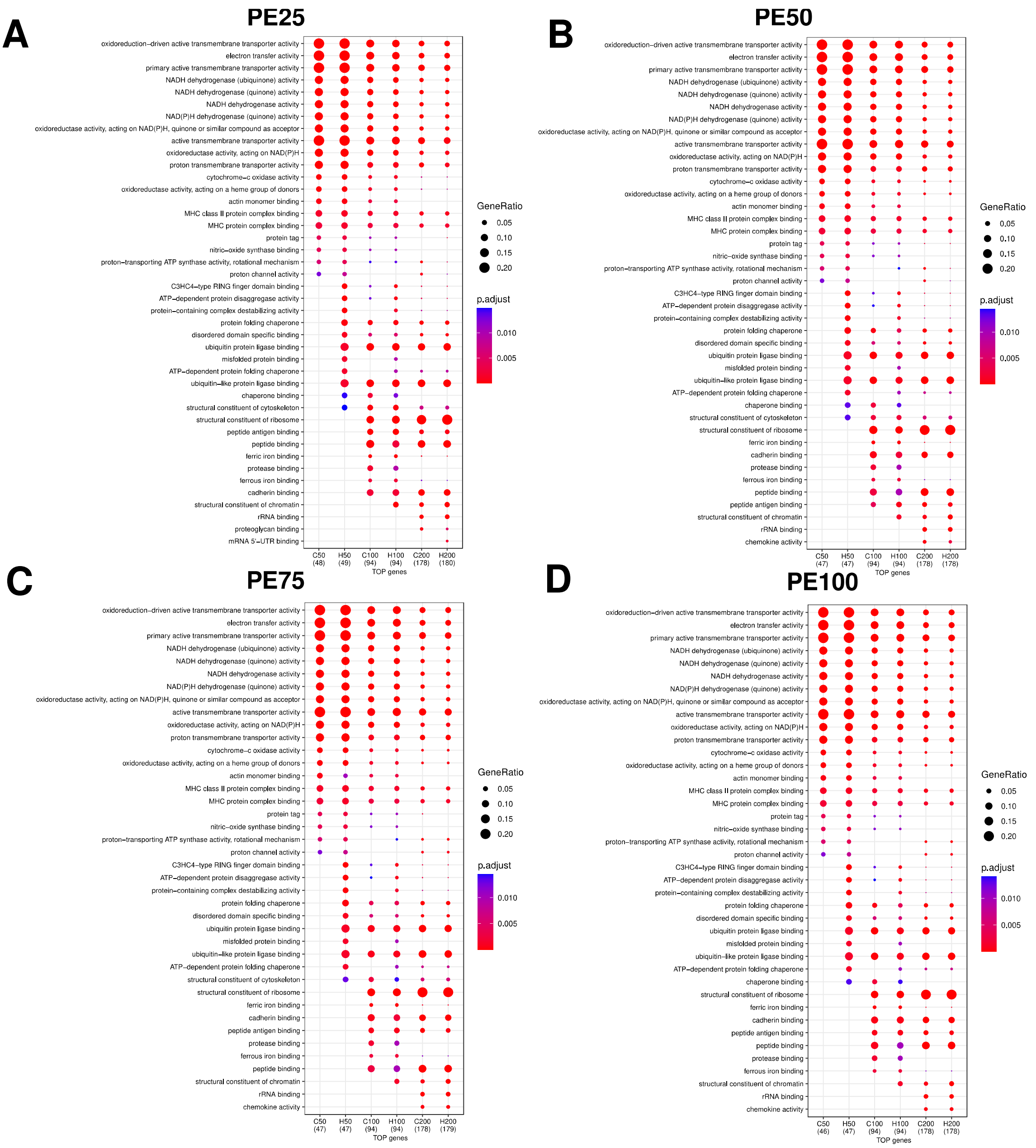


**Additional Figure 6. Functional misrepresentation by HTSeq-count (--nonunique all) for computational trimming of human PE100 RNA-seq library.** Dot plot showing gene ontology (GO) enrichment analysis of the 50, 100, and 200 protein-coding genes with the highest expression values as computed by HTSeq-count using parameter ("--nonunique all") ("H50", "H100", and "H200", respectively) or our "multimapper-aware" strategy ("C50", "C100", and "C200", respectively). **(A)** PE25. **(B)** PE50. **(C)** PE75. **(D)** PE100. For complete description, see legend of Figure 2C.

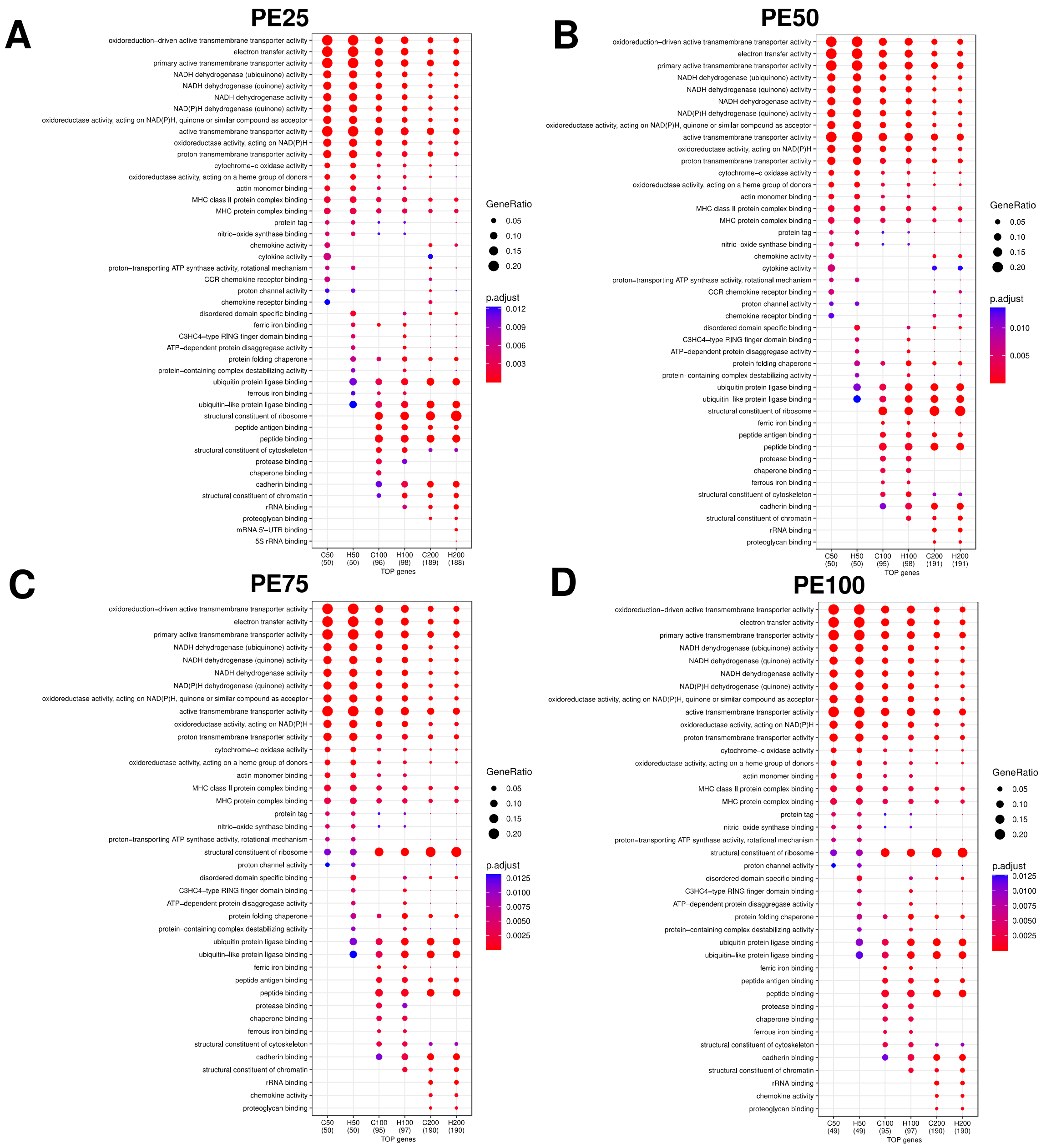


**Additional Figure 7. Functional misrepresentation by HTSeq-count (--nonunique fraction) for computational trimming of human PE100 RNA-seq library.** Dot plot showing gene ontology (GO) enrichment analysis of the 50, 100, and 200 protein-coding genes with the highest expression values as computed by HTSeq-count using parameter ("--nonunique fraction") ("H50", "H100", and "H200", respectively) or our "multimapper-aware" strategy ("C50", "C100", and "C200", respectively). **(A)** PE25. **(B)** PE50. **(C)** PE75. **(D)** PE100. For complete description, see legend of Figure 2C.

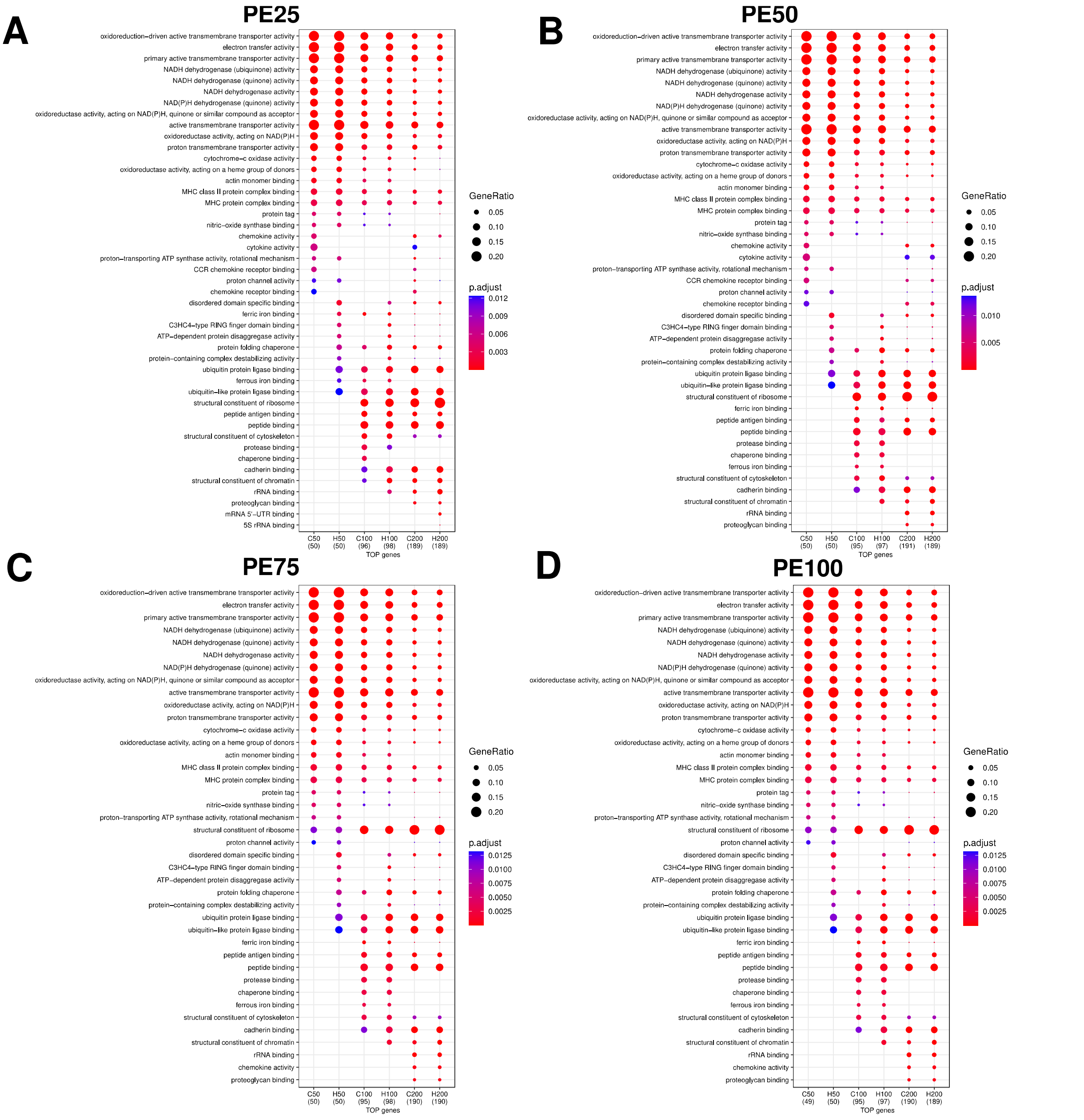


**Additional Figure 8. Functional misrepresentation by HTSeq-count (--nonunique random) for computational trimming of human PE100 RNA-seq library.** Dot plot showing gene ontology (GO) enrichment analysis of the 50, 100, and 200 protein-coding genes with the highest expression values as computed by HTSeq-count using parameter (“--nonunique random”) (“H50”, “H100”, and “H200”, respectively) or our “multimapper-aware” strategy (“C50”, “C100”, and “C200”, respectively). **(A)** PE25. **(B)** PE50. **(C)** PE75. **(D)** PE100. For complete description, see legend of Figure 2C.

The parameter “--nonunique” of HTSeq-count has different modes (“none” (default), “all”, “fraction”, “random”) to decide how to handle fragments assigned to more than one gene in the overlap. For a sanity check, we filtered out overlapping genes and genes from the mitochondrial chromosome. Indeed, we still observed misquantification of genes by HTSeq-count (**Additional Figures 9-12**), suggesting that the gene misquantifications come from the multimappers.

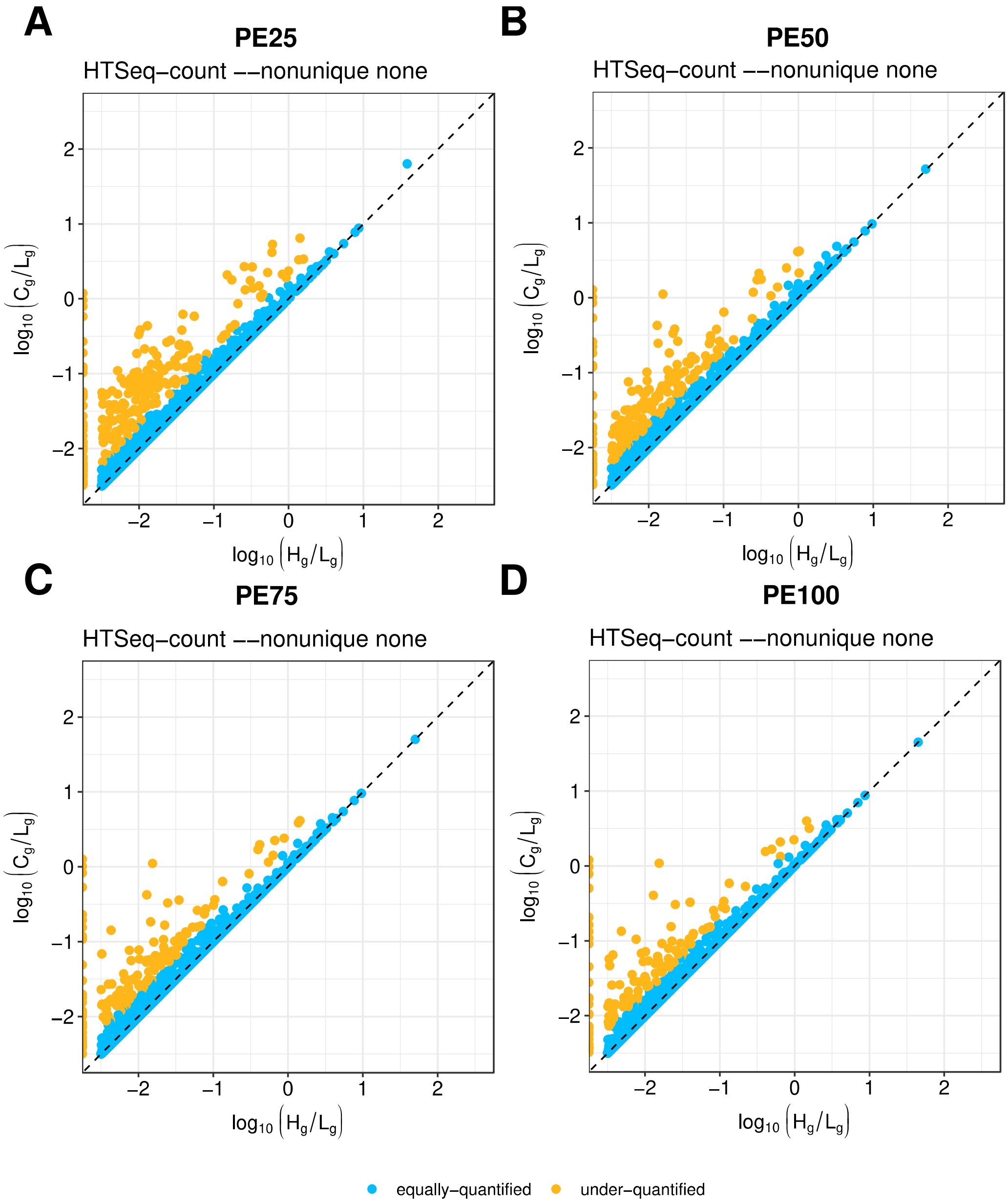


**Additional Figure 9. Gene misquantification by HTSeq-count (--nonunique none) for computational trimming of human PE100 RNA-seq library without overlapping and mitochondrial genes.** Scatter plot showing under-quantified protein-coding genes by HTSeq-count using default parameters (“--nonunique none”; x-axis), when comparing to “multimapper-aware” expression values (y-axis). Percentage of expressed genes that are under-quantified when discarding multimappers for **(A)** PE25: 13% (1,181 out of 8,967). **(B)** PE50: 8% (715 out of 8,632). **(C)** PE75: 6.7% (577 out of 8,572). **(D)** PE100: 6% (516 out of 8,508). For complete description, see legend of Figure 2B.

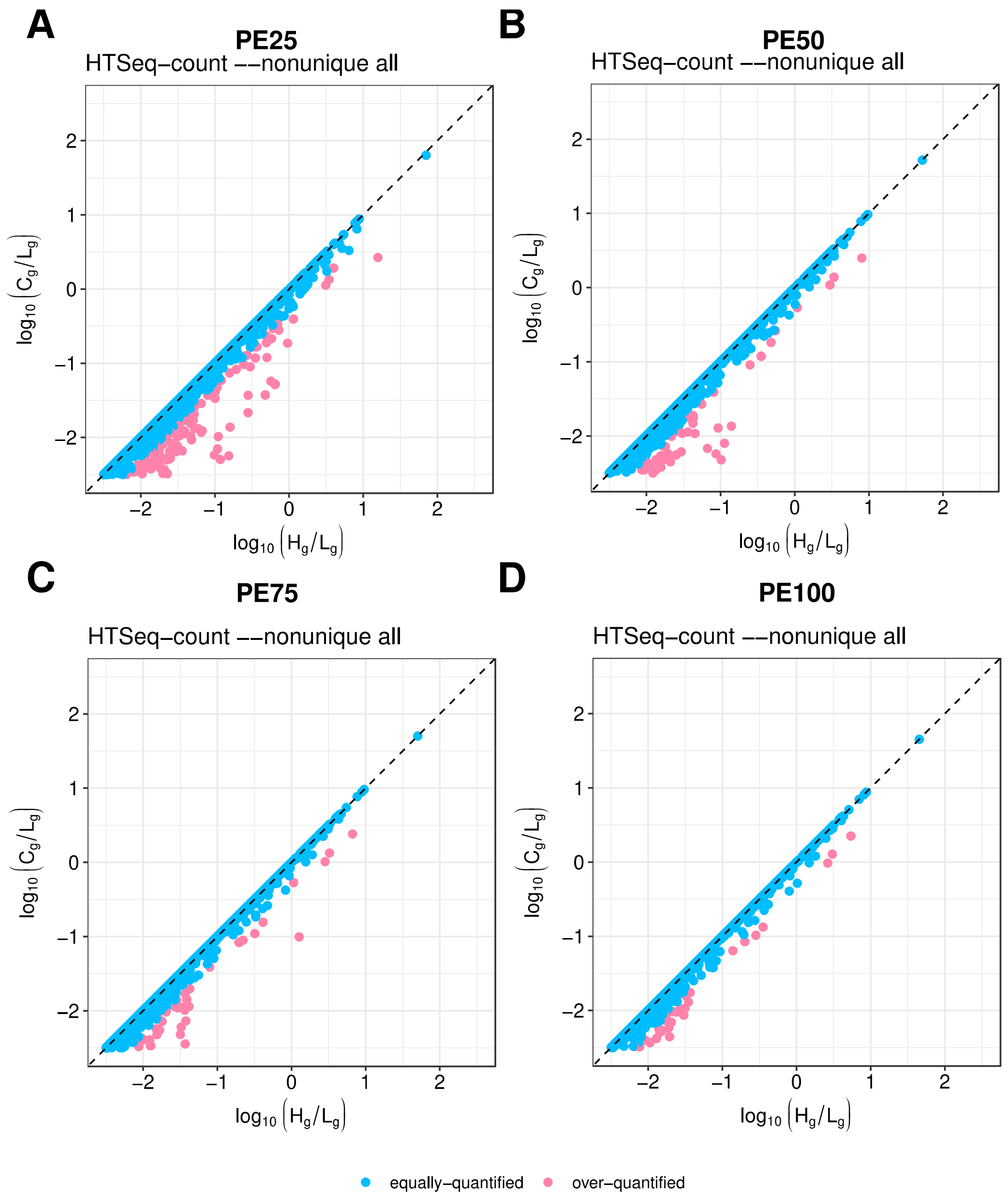


**Additional Figure 10. Gene misquantification by HTSeq-count (--nonunique all) for computational trimming of human PE100 RNA-seq library without overlapping and mitochondrial genes.** Scatter plot showing under-quantified protein-coding genes by HTSeq-count using parameter ("--nonunique all"; x-axis), when comparing to "multimapper-aware" expression values (y-axis). Percentage of expressed genes that are over-quantified when discarding multimappers for **(A)** PE25: 5% (445 out of 9,086). **(B)** PE50: 2.6% (228 out of 8,690). **(C)** PE75: 2% (172 out of 8,627). **(D)** PE100: 1.6% (137 out of 8,558). For complete description, see legend of Figure 2B.

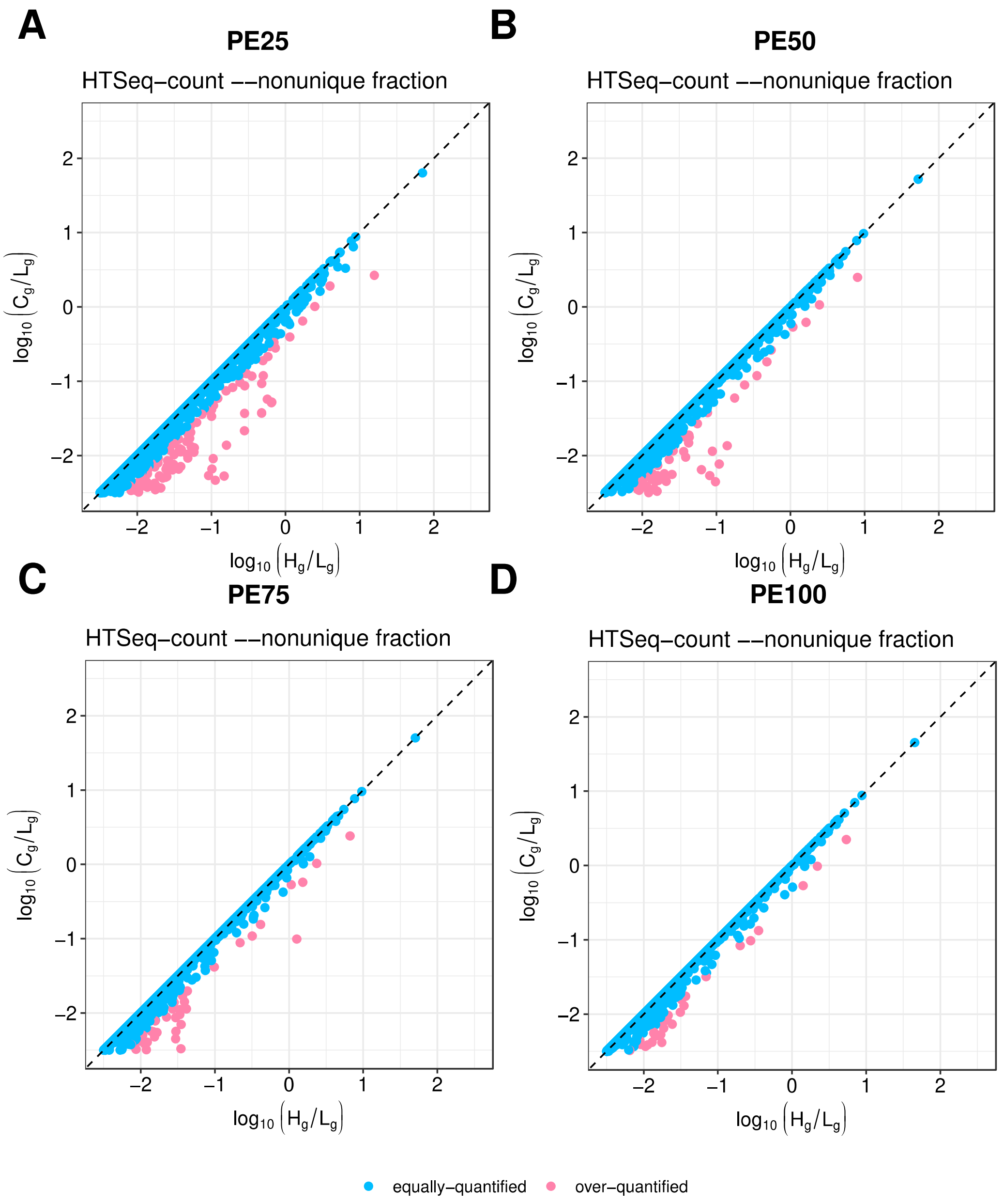


**Additional Figure 11. Gene misquantification by HTSeq-count (--nonunique fraction) for computational trimming of human PE100 RNA-seq library without overlapping and mitochondrial genes.** Scatter plot showing under-quantified protein-coding genes by HTSeq-count using parameter ("--nonunique fraction"; x-axis), when comparing to "multimapper-aware" expression values (y-axis). Percentage of expressed genes that are over-quantified when discarding multimappers for **(A)** PE25: 5% (433 out of 9,086). **(B)** PE50: 2.6% (223 out of 8,690). **(C)** PE75: 2% (167 out of 8,627). **(D)** PE100: 1.6% (134 out of 8,558). For complete description, see legend of Figure 2B.

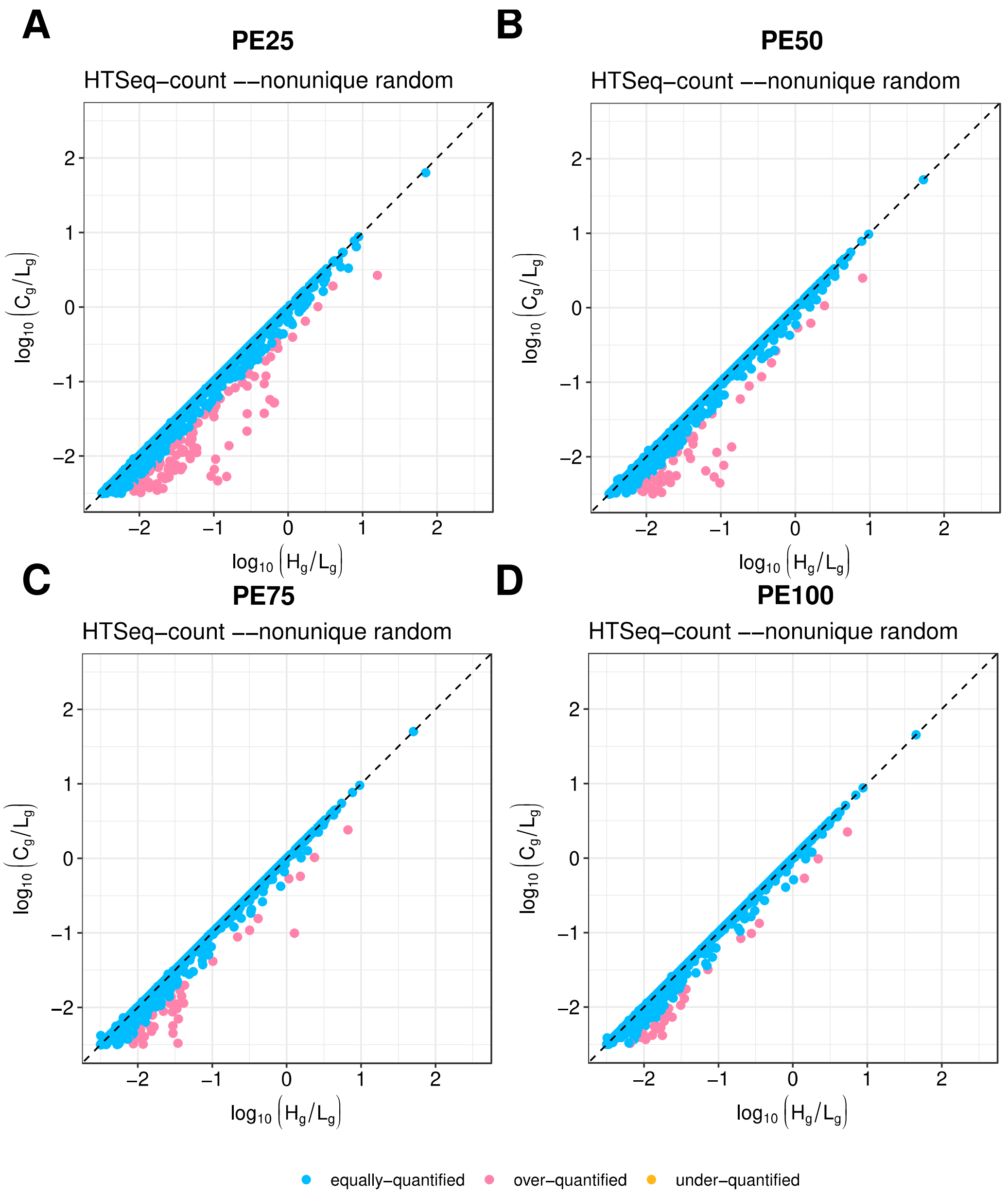


**Additional Figure 12. Gene misquantification by HTSeq-count (--nonunique random) for computational trimming of human PE100 RNA-seq library without overlapping and mitochondrial genes.** Scatter plot showing under-quantified protein-coding genes by HTSeq-count using parameter ("--nonunique random"; x-axis), when comparing to "multimapper-aware" expression values (y-axis). Percentage of expressed genes for **(A)** PE25: 5% (453 out of 9,086) are over-quantified and 0.5% (45 out of 9,086) are under-quantified. **(B)** PE50: 2.6% (224 out of 8,690) are over-quantified and 0.18% (16 out of 8,690) are under-quantified. **(C)** PE75: 2% (171 out of 8,627) are over-quantified and 0.22% (19 out of 8,627) are under-quantified. **(D)** PE100: 1.6% (137 out of 8,558) are over-quantified and 0.16% (14 out of 8,558) are under-quantified. For complete description, see legend of Figure 2B.

Moreover, we still found misrepresented functional GO terms when filtering out overlapping genes and genes from the mitochondrial chromosome (**Additional Figures 13-16**), suggesting that the problem comes, indeed, from multimappers.

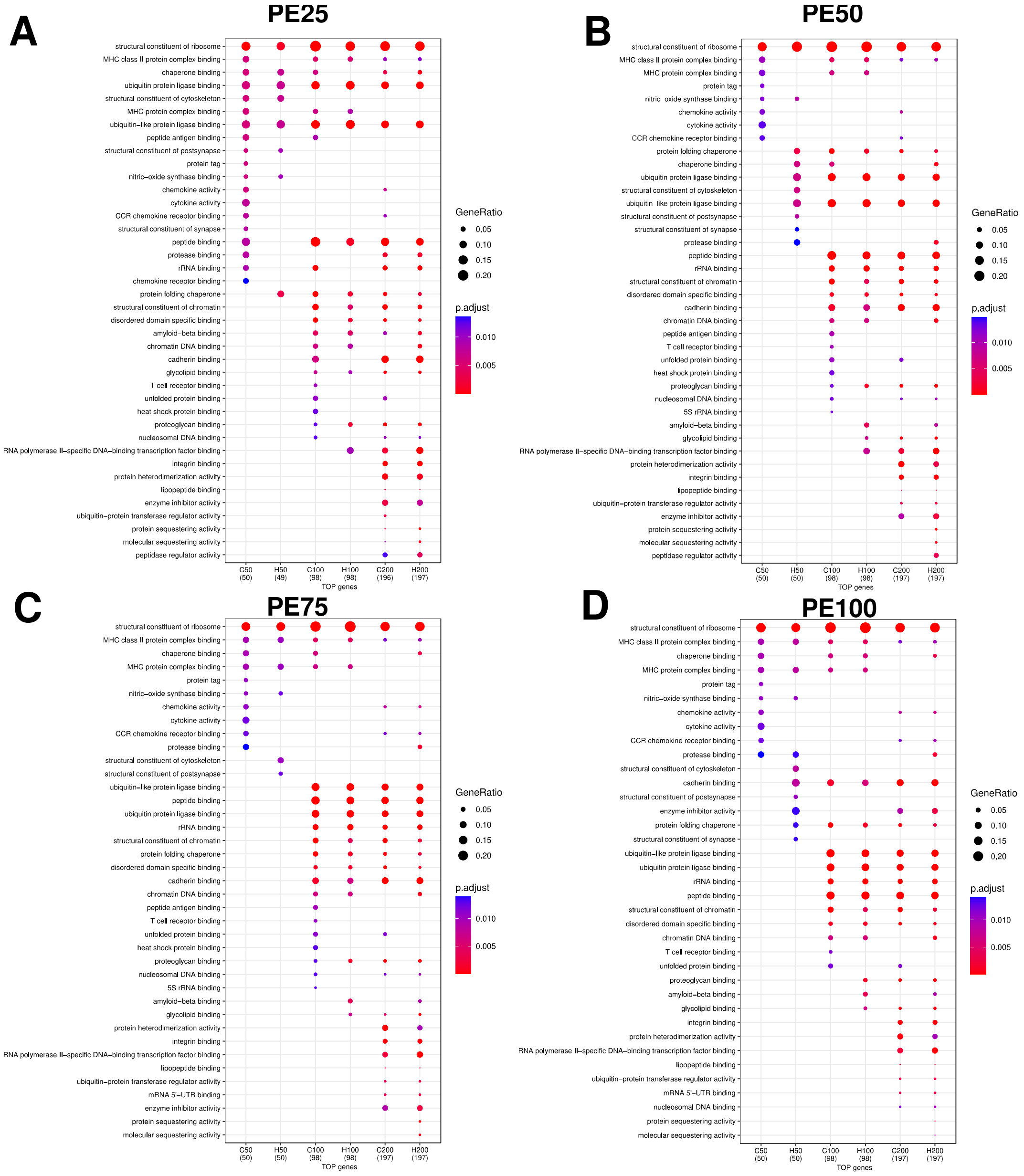


**Additional Figure 13. Functional misrepresentation by HTSeq-count (--nonunique none) for computational trimming of human PE100 RNA-seq library without overlapping and mitochondrial genes.** Dot plot showing gene ontology (GO) enrichment analysis of the 50, 100, and 200 protein-coding genes with the highest expression values as computed by HTSeq-count using default parameters ("--nonunique none") ("H50", "H100", and "H200", respectively) or our "multimapper-aware" strategy ("C50", "C100", and "C200", respectively). **(A)** PE25. **(B)** PE50. **(C)** PE75. **(D)** PE100. For complete description, see legend of Figure 2C.

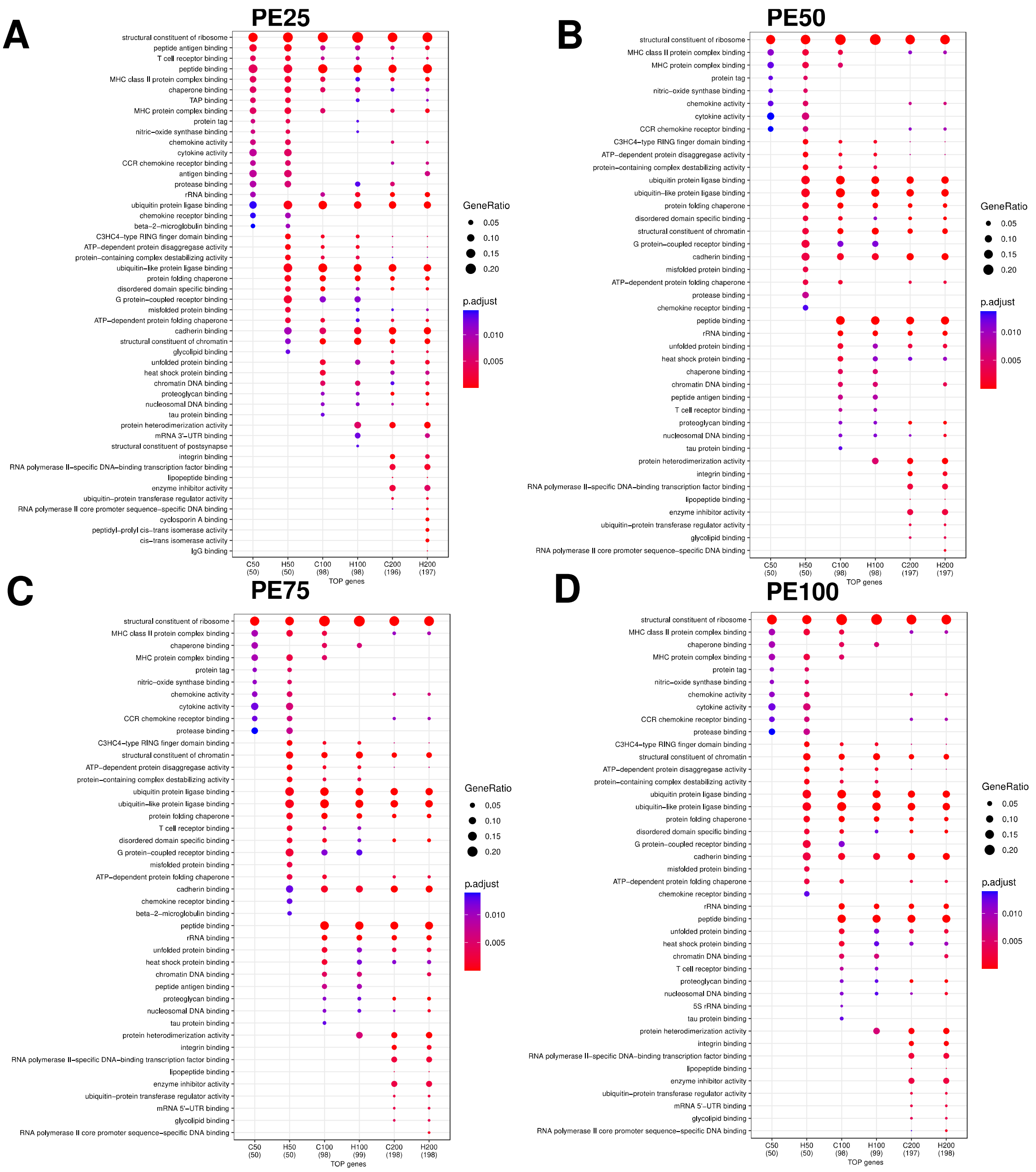


**Additional Figure 14. Functional misrepresentation by HTSeq-count (--nonunique all) for computational trimming of human PE100 RNA-seq library without overlapping and mitochondrial genes.** Dot plot showing gene ontology (GO) enrichment analysis of the 50, 100, and 200 protein-coding genes with the highest expression values as computed by HTSeq-count using default parameters ("--nonunique all") ("H50", "H100", and "H200", respectively) or our "multimapper-aware" strategy ("C50", "C100", and "C200", respectively). **(A)** PE25. **(B)** PE50. **(C)** PE75. **(D)** PE100. For complete description, see legend of Figure 2C.

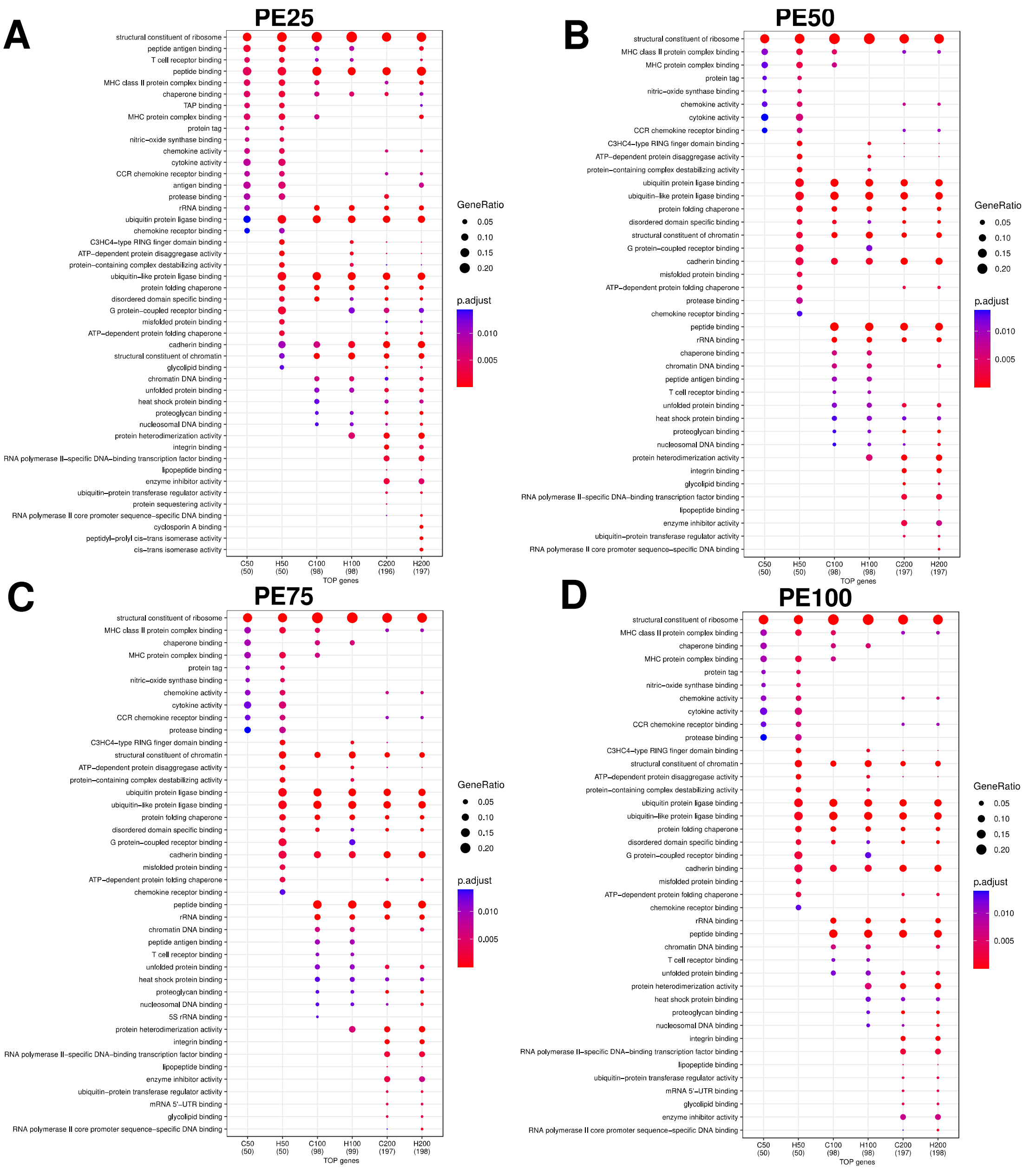


**Additional Figure 15. Functional misrepresentation by HTSeq-count (--nonunique fraction) for computational trimming of human PE100 RNA-seq library without overlapping and mitochondrial genes.** Dot plot showing gene ontology (GO) enrichment analysis of the 50, 100, and 200 protein-coding genes with the highest expression values as computed by HTSeq-count using default parameters ("--nonunique fraction") ("H50", "H100", and "H200", respectively) or our "multimapper-aware" strategy ("C50", "C100", and "C200", respectively). **(A)** PE25. **(B)** PE50. **(C)** PE75. **(D)** PE100. For complete description, see legend of Figure 2C.

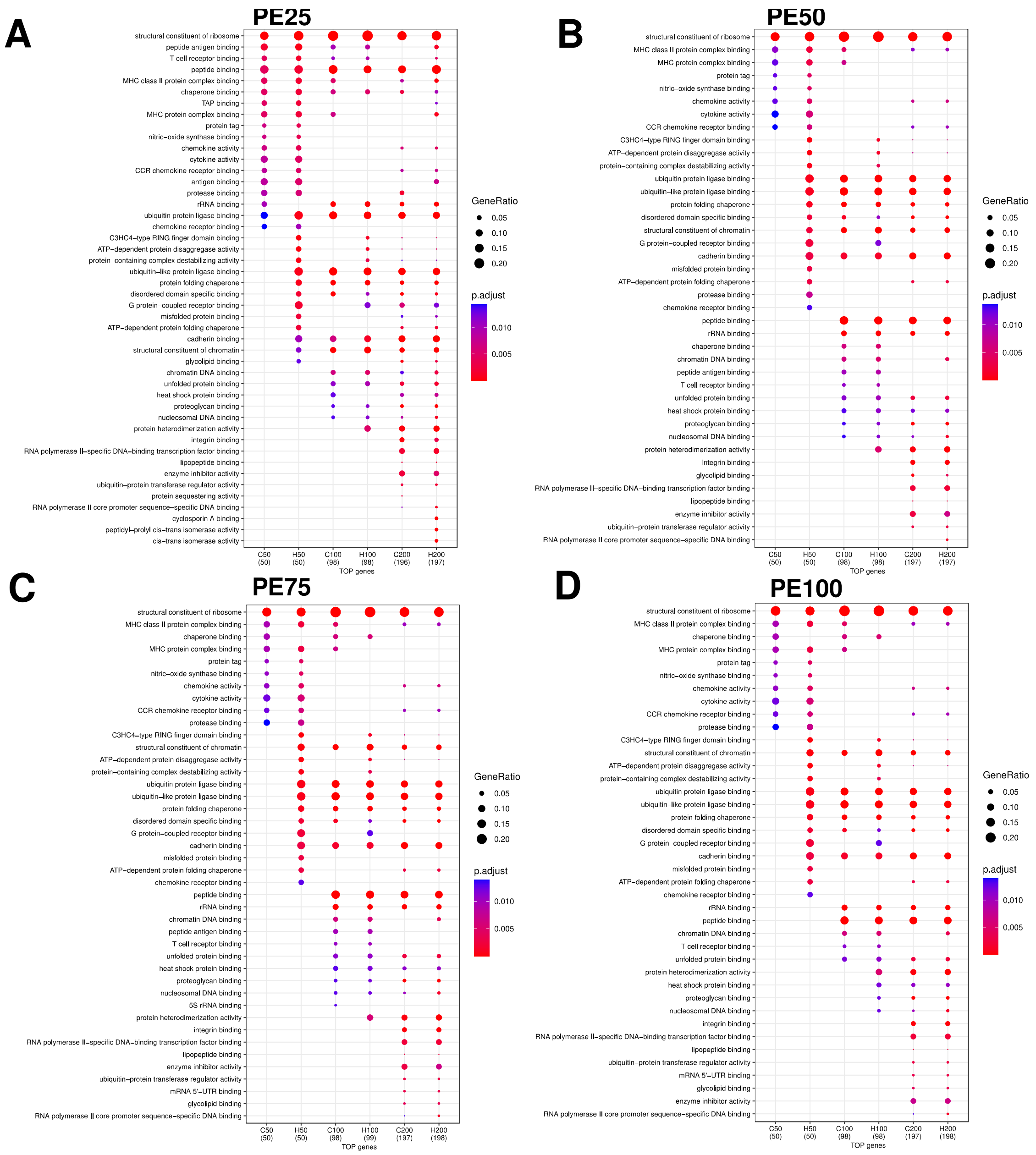


**Additional Figure 16. Functional misrepresentation by HTSeq-count (--nonunique random) for computational trimming of human PE100 RNA-seq library without overlapping and mitochondrial genes.** Dot plot showing gene ontology (GO) enrichment analysis of the 50, 100, and 200 protein-coding genes with the highest expression values as computed by HTSeq-count using default parameters (“--nonunique random”) (“H50”, “H100”, and “H200”, respectively) or our “multimapper-aware” strategy (“C50”, “C100”, and “C200”, respectively). **(A)** PE25. **(B)** PE50. **(C)** PE75. **(D)** PE100. For complete description, see legend of Figure 2C.

For the mouse RNA-seq library ("PE75", see Methods), we saw that about 4% (468 out of 12,561) of the expressed genes were under-quantified by HTSeq-count using default parameters ("--nonunique none"). As a mean of verification, we evaluated other HTSeq-count "--nonunique" options (all, fraction and random) (**Additional Figure 17**). For the modes all (**Additional Figure 17A**) and fraction (**Additional Figure 17B**), the percentage of over-quantified genes was ~2%. For the "--nonunique random", about 2% of the expressed genes were over-quantified, and about 0.5% were under-quantified (**Additional Figure 17C**).

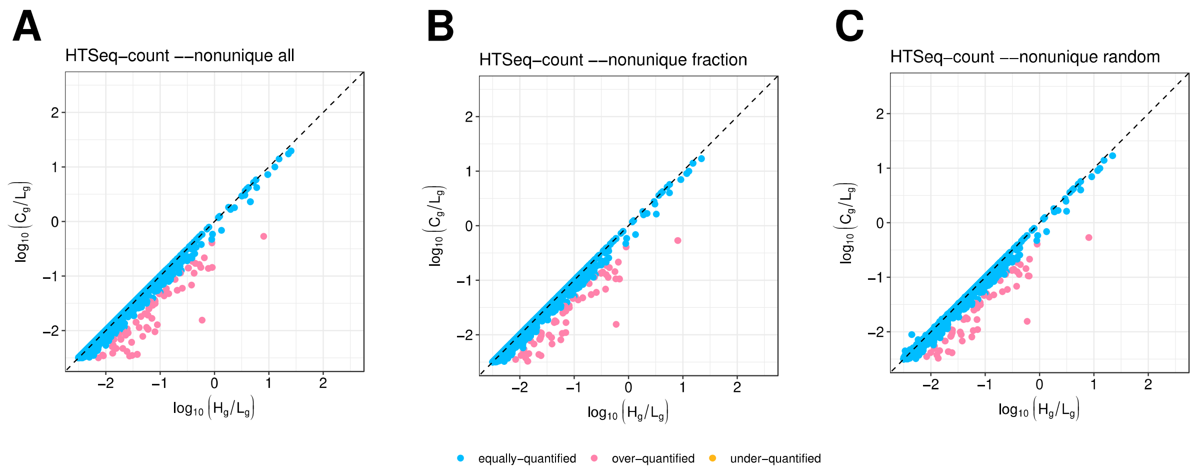


**Additional Figure 17. Gene misquantification by HTSeq-count for mouse PE75 RNA-seq library.** Scatter plot showing under- and over-quantified protein-coding genes by HTSeq-count using different ("--nonunique") parameter (x-axis), when comparing to "multimapper-aware" expression values (y-axis). Percentage of expressed genes for **(A)** HTSeq-count "--nonunique all": 2% (289 out of 12,803) are over-quantified. **(B)** HTSeq-count "--nonunique fraction": 2% (282 out of 12,803) are over-quantified. **(C)** HTSeq-count "--nonunique random": 2% (291 out of 12,803) are over-quantified and 0.5% (59 out of 12,803) are under-quantified. For complete description, see legend of Figure 2B.

In addition, we evaluated other HTSeq-count “--nonunique” options (all, fraction and random) and found other misrepresented functional GO terms (**Additional Figure 18**).

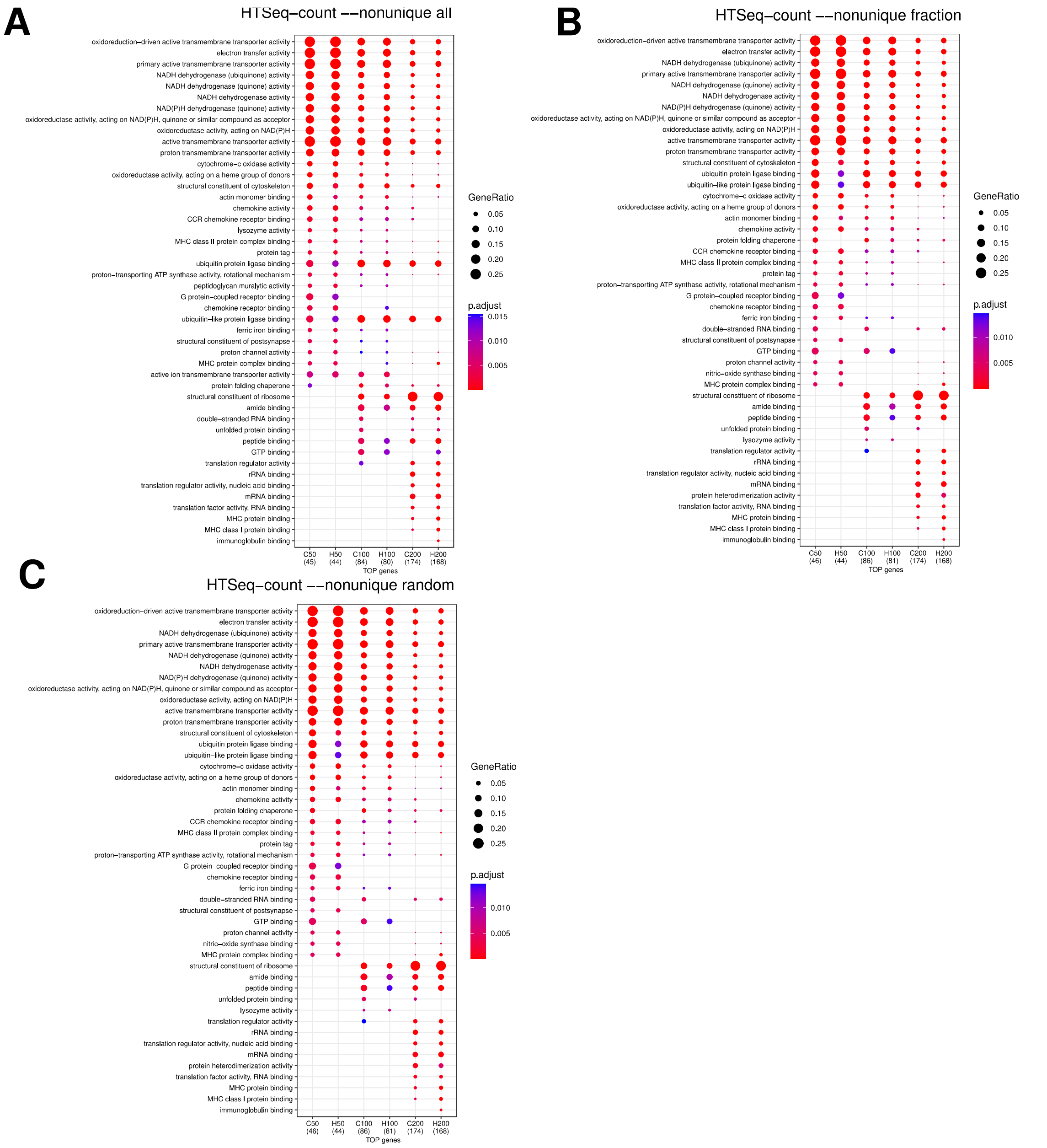


**Additional Figure 18. Functional misrepresentation by HTSeq-count for mouse PE75 RNA-seq library.** Dot plot showing gene ontology (GO) enrichment analysis of the 50, 100, and 200 protein-coding genes with the highest expression values as computed by HTSeq-count using different (“--nonunique”) parameter (“H50”, “H100”, and “H200”, respectively) or our “multimapper-aware” strategy (“C50”, “C100”, and “C200”, respectively). **(A)** HTSeq-count “--nonunique all”. **(B)** HTSeq-count “--nonunique fraction”. **(C)** HTSeq-count “--nonunique random”. For complete description, see legend of Figure 2C.

For a sanity check in the mouse RNA-seq library, we also filtered out overlapping genes and genes from the mitochondrial chromosome. Likewise to human RNA-seq libraries, we observed misquantification of genes by HTSeq-count (**Additional Figure 19**), suggesting that the gene misquantifications come from the multimappers.

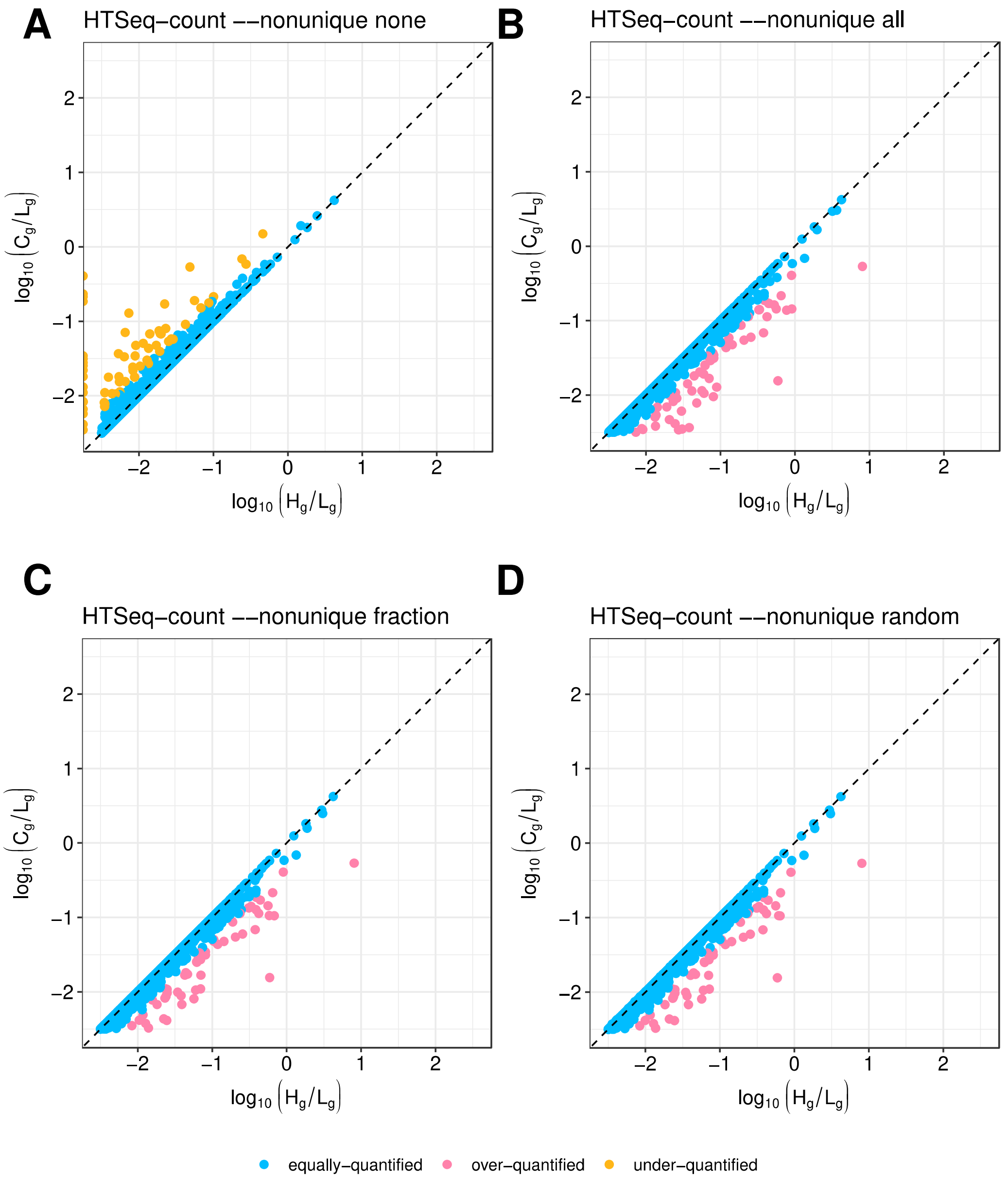


**Additional Figure 19. Gene misquantification by HTSeq-count for mouse PE75 RNA-seq library without overlapping and mitochondrial genes.** Scatter plot showing under- and over-quantified protein-coding genes by HTSeq-count using different ("--nonunique") parameter (x-axis), when comparing to "multimapper-aware" expression values (y-axis). Percentage of expressed genes for **(A)** HTSeq-count "--nonunique none": 4% (379 out of 9,477) are under-quantified. **(B)** HTSeq-count "--nonunique all": 2.7% (254 out of 9,513) are over-quantified. **(C)** HTSeq-count "--nonunique fraction": 2.6% (248 out of 9,513) are over-quantified. **(D)** HTSeq-count "--nonunique random": 2.6% (245 out of 9,513) are over-quantified and 0.14% (13 out of 9,513) are under-quantified. For complete description, see legend of Figure 2B.

Finally, we performed functional analysis for mouse RNA-seq library and we found misrepresented functional GO terms when filtering out overlapping genes and genes from the mitochondrial chromosome (**Additional Figure 20**), reinforcing that multimappers lead to the observed functional misrepresentation.

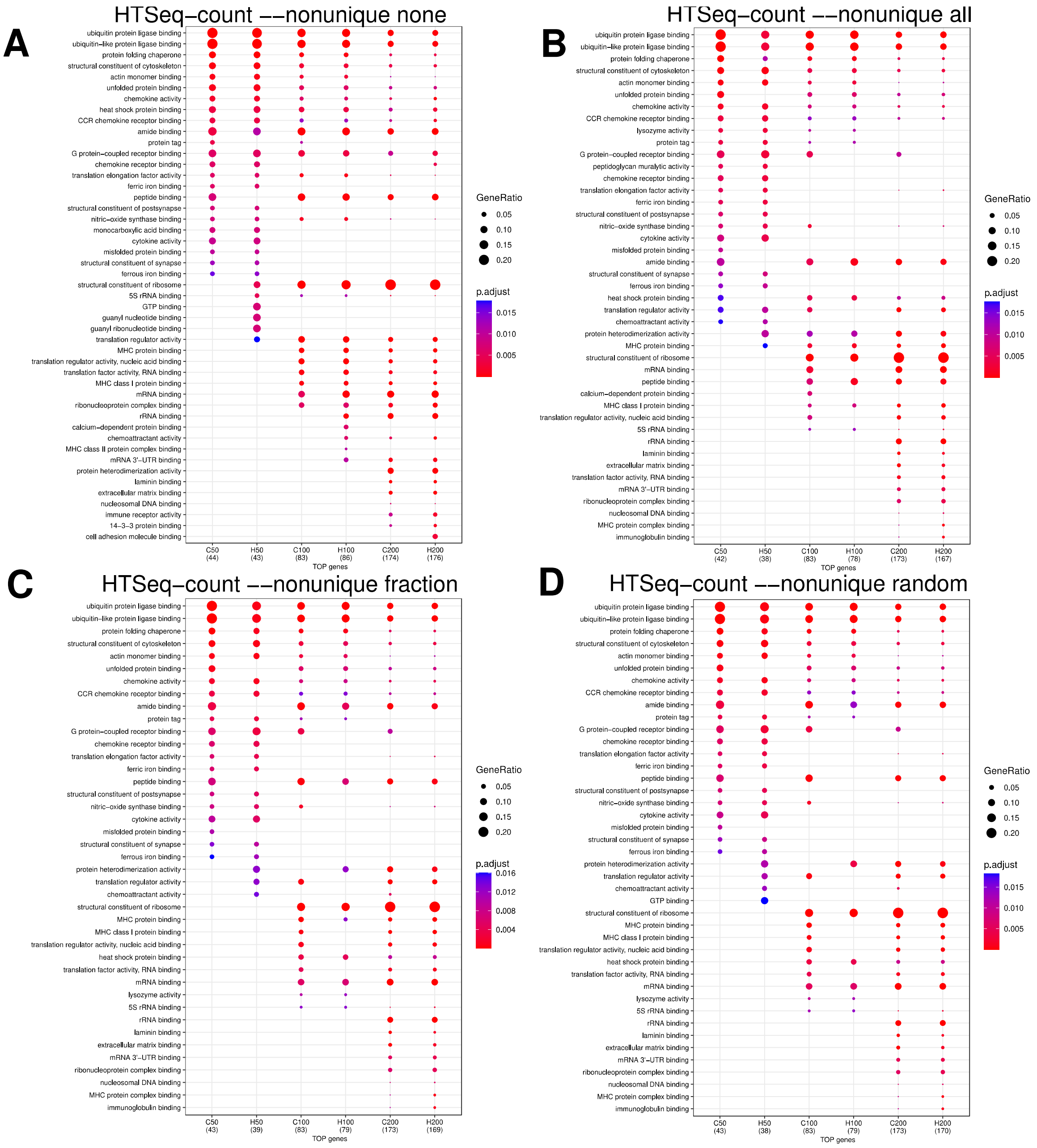


**Additional Figure 20. Functional misrepresentation by HTSeq-count for mouse PE75 RNA-seq library without overlapping and mitochondrial genes.** Dot plot showing gene ontology (GO) enrichment analysis of the 50, 100, and 200 protein-coding genes with the highest expression values as computed by HTSeq-count using different ("--nonunique") parameter ("H50", "H100", and "H200", respectively) or our "multimapper-aware" strategy ("C50", "C100", and "C200", respectively). **(A)** HTSeq-count "--nonunique none". **(B)** HTSeq-count "--nonunique all". **(C)** HTSeq-count "--nonunique fraction". **(D)** HTSeq-count "--nonunique random". For complete description, see legend of Figure 2C.
